# Intra-helical salt bridge contribution to membrane protein insertion

**DOI:** 10.1101/2021.02.24.432724

**Authors:** Gerard Duart, John Lamb, Juan Ortiz-Mateu, Arne Elofsson, Ismael Mingarro

## Abstract

Salt bridges between negatively (D, E) and positively charged (K, R, H) amino acids play an important role in protein stabilization. This has a more prevalent effect in membrane proteins where polar amino acids are exposed to a very hydrophobic environment. In transmembrane (TM) helices the presence of charged residues can hinder the insertion of the helices into the membrane. This can sometimes be avoided by TM region rearrangements after insertion, but it is also possible that the formation of salt bridges could decrease the cost of membrane integration. However, the presence of intra-helical salt bridges in TM domains and their effect on insertion has not been properly studied yet. In this work, we use an analytical pipeline to study the prevalence of charged pairs of amino acid residues in TM α-helices, which shows that potentially salt-bridge forming pairs are statistically over-represented. We then selected some candidates to experimentally determine the contribution of these electrostatic interactions to the translocon-assisted membrane insertion process. Using both *in vitro* and *in vivo* systems, we confirm the presence of intra-helical salt bridges in TM segments during biogenesis and determined that they contribute between 0.5-0.7 kcal/mol to the apparent free energy of membrane insertion (ΔG_app_). Our observations suggest that salt bridge interactions can be stabilized during translocon-mediated insertion and thus could be relevant to consider for the future development of membrane protein prediction software.

## INTRODUCTION

Most integral membrane proteins have to insert their transmembrane (TM) segments into the lipid bilayer in a helical conformation and then acquire a defined three-dimensional structure by packaging their helices (Martínez-Gil et al., 2011). α-Helical TM segments are largely composed of apolar residues because of the hydrophobic nature of the membrane environment. Nevertheless, in some cases, it is necessary for the protein activity to include polar amino acids in a TM region in order to develop a functional or structural role (Baeza-Delgado et al., 2012). This fact is sometimes not contemplated in modern membrane topology prediction tools (Tsirigos et al., 2018; 2015), in which the presence of charged amino acids in a sequence automatically suppose a penalty increase in the predicted free energy (ΔG_pred_) of insertion. The presence of polar amino acids in TM regions is more frequent than what would be expected (Bañó-Polo et al., 2012), especially when these are in pairs on the same face of an α-helix.

Salt bridges are electrostatic interactions between negatively (D, E) and positively charged (K, R, H) amino acids that play an important role in protein stabilization (Marqusee and Baldwin, 1987). Many studies have shown that pairs of charged residues that form potential salt-bridges stabilize soluble α-helices (Donald et al., 2011). Salt bridges play an important role in the folding of globular proteins and, despite their low occurrence in TM domains, it seems that the contribution in membrane protein stability could be even more determinant. This contribution is especially important in membrane protein biogenesis, as salt bridges help to bury the polarity of charged residues in a hydrophobic environment (Mbaye et al., 2019). Apart from that, it has been suggested that potential salt bridges could help in the insertion of TM α-helices (Baeza-Delgado et al., 2016; Bañó-Polo et al., 2012), even though their predicted ΔG penalty is well above what is usually seen for TM segments.

To investigate the potential formation of intra-helical salt bridges in TM α-helices, we analyzed the composition of the TM domains from membrane proteins of known structures looking for preferences in the pairing of charged amino acids. This analysis showed that charged residue pairing is more prevalent than expected for pairs located on the same face of α-helices within membranes. Likely, salt bridge formation on the same face of α-helices reduces the unfavorable energetics of inserting ionizable residues into the hydrophobic membrane core (Chin and Heijne, 2000). We use this knowledge together with the known membrane protein structures to generate a list of potential candidates for further *in vitro* and *in vivo* experiments.

In this work, we have studied the presence of intra-helical salt bridges in TM domains using *in silico*, *in vitro* and *in vivo* systems, showing that despite this being an unexpected phenomenon in nature, salt bridges can be crucial for membrane insertion. Also, we determined in a quantitative manner that the apparent free energy (ΔG_app_) of membrane insertion through the translocon machinery can be decreased between 0.5-0.7 kcal/mol by position-specific charge pair interaction, which is not contemplated in the commonly used ΔG predictors. These findings will lead to a better understanding of the insertion mechanism of TM helices and to improve prediction tools that would more accurately be able to model the presence of charged residues in these helices.

## RESULTS

### Charge pair interactions in model transmembrane helices

To test the contribution of potential salt bridges to the translocon-mediated membrane insertion, we used the vehicle protein leader peptidase (Lep) from *Escherichia coli* (Fig. 1a). The Lep protein consists of two TM segments (H1 and H2) connected by a cytoplasmic loop (P1) and a large C-terminal (P2) domain, which inserts into endoplasmic reticulum (ER)-derived rough microsomes with both termini located in the microsomes lumen. The designed TM segments were inserted into the luminal P2 domain and flanked by two acceptor sites (G1 and G2) for N-linked glycosylation. Glycosylation occurs exclusively in the lumen of the ER (or microsomes) because of the location of the oligosaccharyltransferase (OST) active site (a translocon-associated enzyme responsible for the oligosaccharide transfer) (Braunger et al., 2018). In this case, the engineered glycosylation sites can be used as membrane insertion reporters because G1 will always be glycosylated due to its native luminal localization, but G2 will be glycosylated only upon translocation of the analyzed sequence across the microsomal membrane. A singly glycosylated construct in which a tested sequence is inserted into the membrane has a molecular mass ∼2.5 kDa higher than the molecular mass of Lep molecule expressed in the absence of microsomes; the molecular mass shifts by ∼5 kDa upon double glycosylation, which facilitates its identification by gel electrophoresis when expressed in the presence of [^35^S-labeled] amino acids. Then, *in vitro* transcription/translation of these chimeric proteins in the presence of rough microsomal membranes (RMs) allows for accurate and quantitative description of membrane insertion of designed sequences (Bañó-Polo et al., 2019; Hessa et al., 2005; 2007; Tamborero et al., 2011). The degree of membrane insertion is quantified by analyzing the fractions of singly glycosylated (i.e., membrane inserted) and doubly glycosylated (i.e., non-inserted) molecules, which can be expressed as an experimental apparent free energy of membrane insertion, ΔG_exp_ (see Materials and Methods) (Hessa et al., 2005).

**Figure 1.**
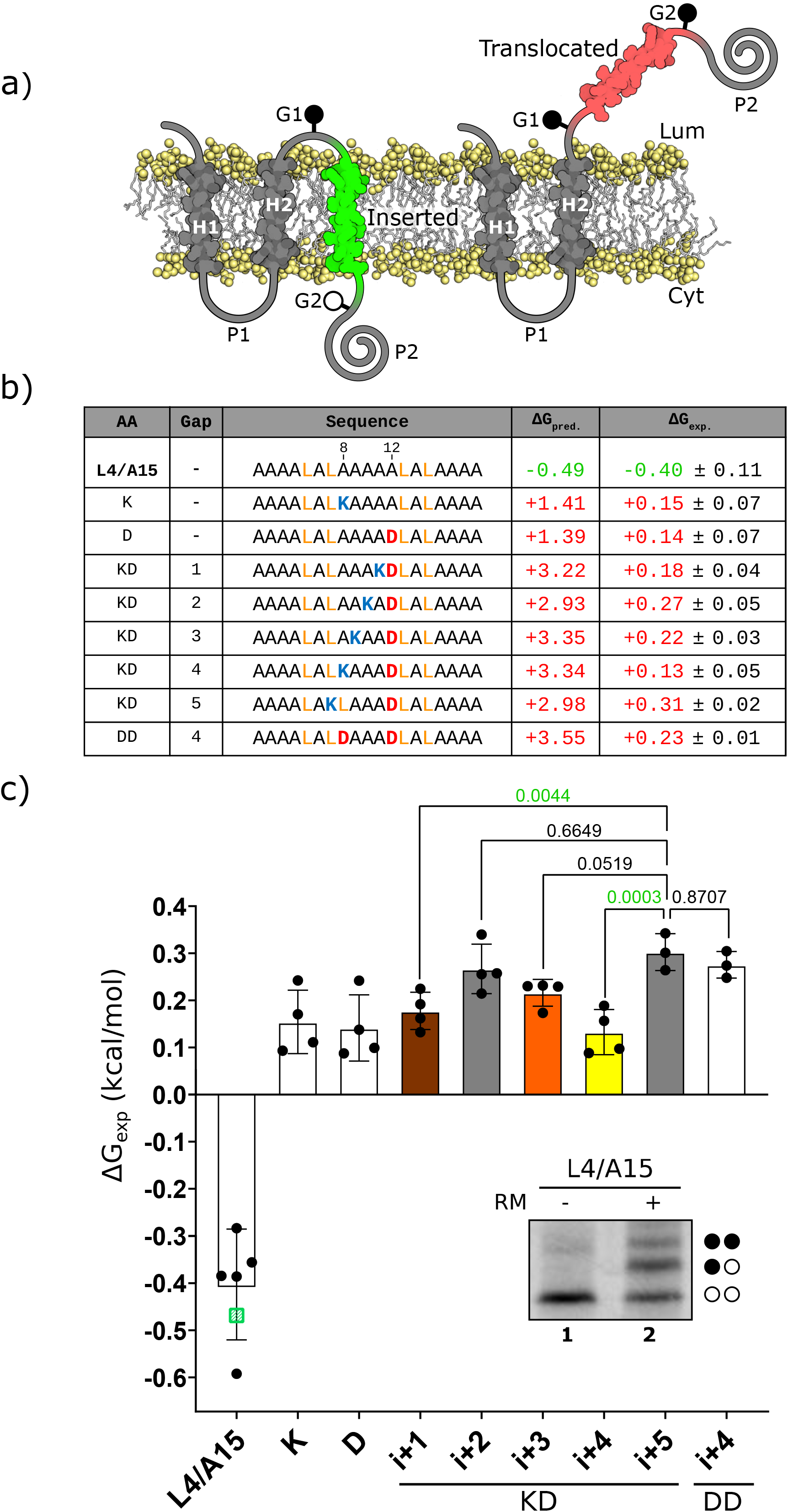
Effects on membrane insertion of single or pairs of Asp and Lys residues in a model TM segment. **(a)** Schematic representation of the leader peptidase (Lep) model protein. G1 and G2 denote artificial glycosylation acceptor sites. The sequence under investigation was introduced in the P2 region after H2. Recognition of the tested sequence as a TM by the translocon machinery (highlighted in green) results in the modification of the G1 site but not G2. The Lep chimera will be double glycosylated if the sequence being tested is not recognized as a TMD and thus translocated into the microsomes lumen (shown in red). **(b)** The tested sequences from L4/A15 model TM (including the charged residues, bold), the gap distance, and the predicted ΔG (ΔG_pred_) values in kcal/mol are shown. Amino acids with positive and negative charge are highlighted in blue (K) and red (D) respectively. **(c)** Experimental ΔG (ΔG_exp_) in kcal/mol of each tested sequence in the Lep-based microsomal assay. The mean and standard deviation of at least 3 independent experiments is represented (*n* values: 4 [from 1 to 7] and 3 [8 and 9]). The individual value of each experiment is represented by a solid dot, *p*-values (ordinary one-way ANOVA test with Dunnett correction) are indicated above the corresponding bars with values <0.005 highlighted in green. In addition, a green square represents the experimental ΔG value for the L4/A15 sequence from an earlier study [12]. The wt and single mutants are shown in white bars. Charges at compatible distances with salt bridge formation (*i*, *i*+1; *i*, *i*+3; and *i*, *i*+4) are shown in brown, orange and yellow, respectively. Not compatible distances with salt bridge formation (*i*, *i*+2; and *i*, *i*+5) are shown in dark gray. The inset shows a representative SDS-PAGE gel for L4/A15 construct. The construct was expressed in rabbit reticulocyte lysed in the presence (+RM) or absence (–RM) of rough microsomes. Bands of non-glycosylated proteins are indicated by a white dot; mono and double glycosylated proteins are indicated by one and two black dots, respectively.

We first compared the effects of oppositely charged Lys and Asp residues on the insertion of a 19-residue-long hydrophobic stretch (L4/A15 scaffold, 4 leucines and 15 alanines), which was designed to insert stably into the RM membranes (Hessa et al., 2005), including charges centered in the TM segment at different positions (Fig. 1b) and “insulated” from the surrounding sequence by N-and C-terminal GGPG-and -GPGG tetrapeptides. Single Lys and Asp residues were placed in positions 8 and 12 respectively, and pairs of Lys-Asp residues were designed to cover positions 7-12 (that is, more than one helical turn). When pairs of charged residues are present, our results showed a tendency to insert more efficiently when pair charges were placed in positions (*i*, *i*+1; *i*, *i*+3; *i*, *i*+4) that are permissive with salt bridge formation (Fig. 1c), actually an effect not observed in the predictions (Fig. 1b). Similar results were obtained on a different Leu/Ala background with a slightly higher insertion efficiency (L5/A14, 5 leucines and 14 alanines), those mutants harbouring charged pairs compatible with salt bridge formation (i.e. *i*, *i*+3; *i*, *i*+4) insert more efficiently than the non-compatible one *i*, *i*+5 (Figure S1). Being the insertion of charged residues a thermodynamically inconvenient phenomenon within the membrane environment, it is expected that the sequence context and the amino acid composition of the TM helix would be determinant for salt bridge formation. Accordingly, we scrutinized a large dataset of membrane proteins of known three-dimensional structures in order to focus on natural salt bridges present within TM segments.

### Charged pairs in transmembrane helices

Alpha helices are a common secondary structure in both globular and membrane proteins. We created two main datasets, TM dataset with helical membrane proteins of known structure from the PDBTM-dataset (Kozma et al., 2013), and GLOB dataset with globular alpha helical proteins selected from the SCOP-database (Andreeva et al., 2013; 2019), see methods for the full creation steps. Table 1 shows a breakdown of alpha-helices, charged residues and salt bridges in the two datasets. What is clear is that long alpha helices (≥ 17 residues) form a larger proportion in the TM dataset than in the globular helices (GLOB) dataset, which is logical as most TM helices need to span through the hydrophobic core of the lipid bilayer of thickness ∼30 Å.

**Table 1.**
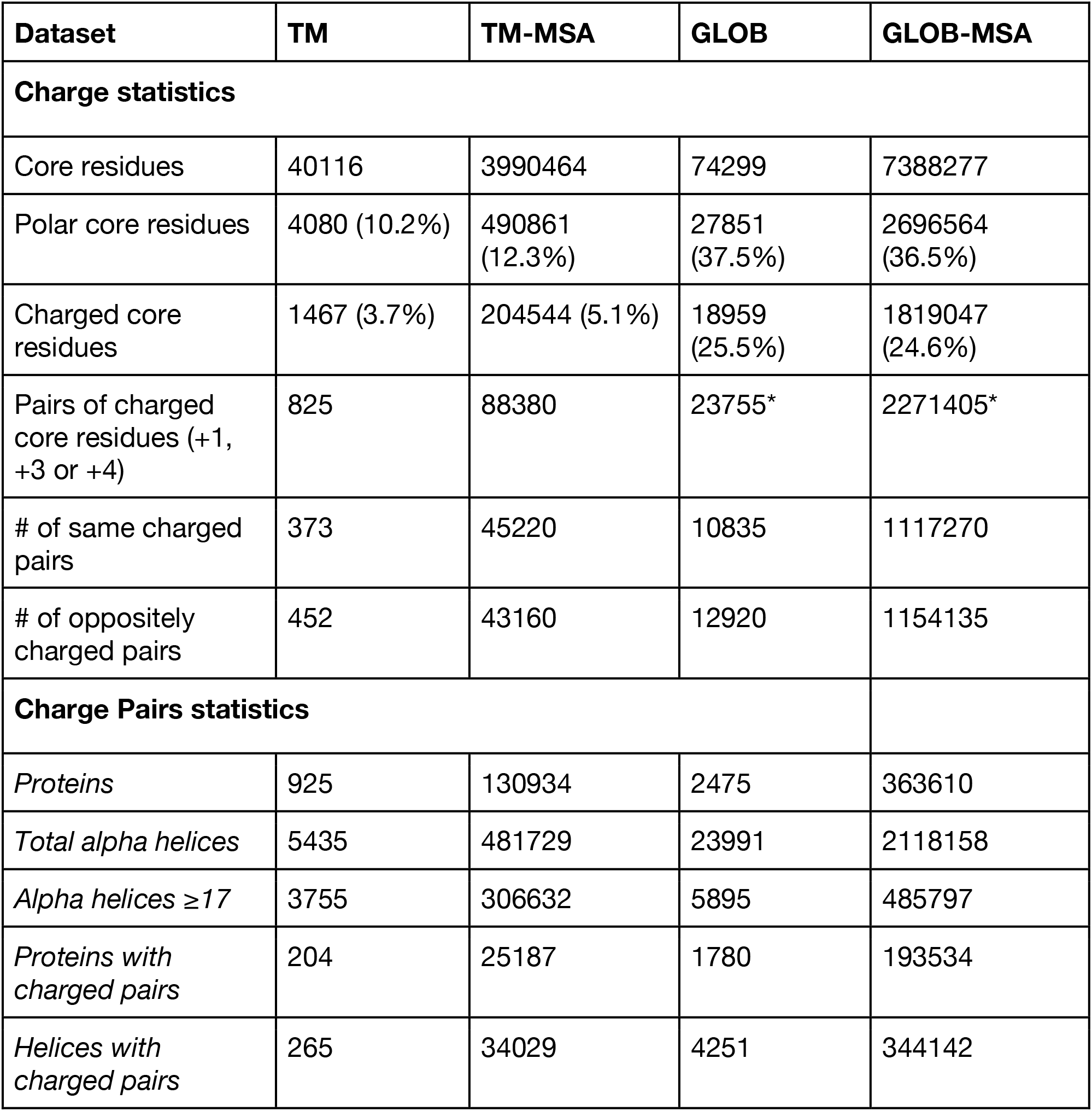
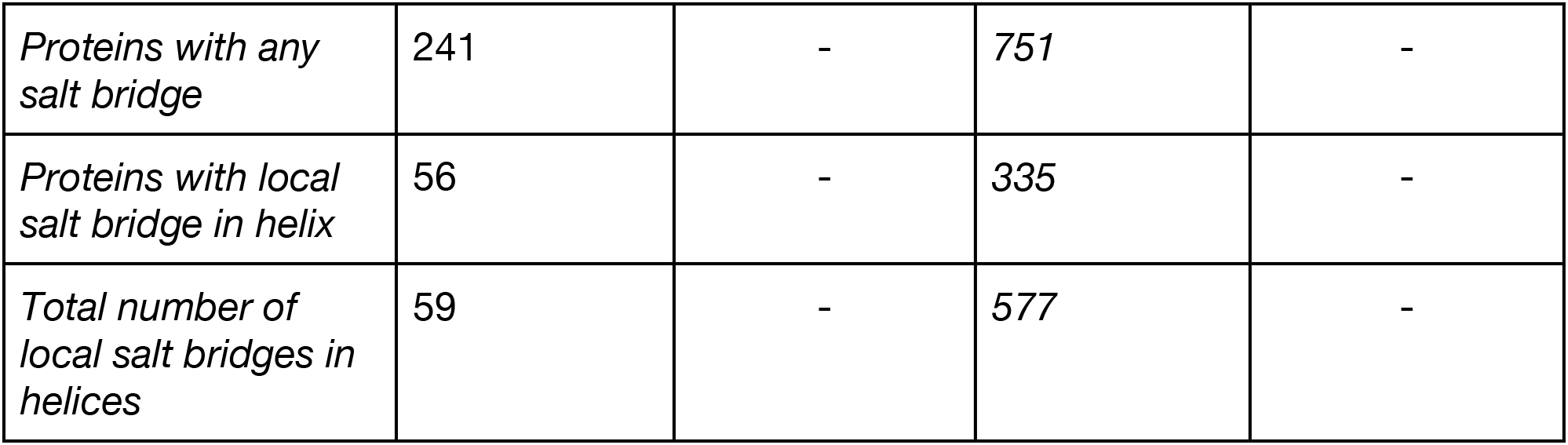
About 10 % of the core residues in the TM dataset are polar and just under 4% are charged residues. This is significantly less than 37.5% and 25.5% respectively in the globular dataset. In the transmembrane dataset, just over half of the charged core residues form a charged pair at distance 1, 3 or 4. *In the case of the globular set, there are more potential charged pairs than charged core residues as multiple residues are counted more than once as they form more than one potential pairing with separation of 1, 3 or 4 steps. It is clearly seen that although the TM dataset contains more helices per protein and has a higher proportion of long (≥ 17 residues) helices it contains significantly fewer charged pairs.

| Dataset | TM | TM-MSA | GLOB | GLOB-MSA |
| --- | --- | --- | --- | --- |
| **Charge statistics** |  |  |  |  |
| Core residues | 40116 | 3990464 | 74299 | 7388277 |
| Polar core residues | 4080 (10.2%) | 490861<br>(12.3%) | 27851<br>(37.5%) | 2696564<br>(36.5%) |
| Charged core residues | 1467 (3.7%) | 204544 (5.1%) | 18959<br>(25.5%) | 1819047<br>(24.6%) |
| Pairs of charged core residues (+1, +3 or +4) | 825 | 88380 | 23755* | 2271405* |
| # of same charged pairs | 373 | 45220 | 10835 | 1117270 |
| # of oppositely charged pairs | 452 | 43160 | 12920 | 1154135 |
| **Charge Pairs statistics** |  |  |  |  |
| *Proteins* | 925 | 130934 | 2475 | 363610 |
| *Total alpha helices* | 5435 | 481729 | 23991 | 2118158 |
| *Alpha helices $\geq 17$* | 3755 | 306632 | 5895 | 485797 |
| *Proteins with charged pairs* | 204 | 25187 | 1780 | 193534 |
| *Helices with charged pairs* | 265 | 34029 | 4251 | 344142 |
| *Proteins with any salt bridge* | 241 | - | 751 | - |
| *Proteins with local salt bridge in helix* | 56 | - | 335 | - |
| *Total number of local salt bridges in helices* | 59 | - | 577 | - |

Our TM dataset showed the same distribution of polar charges as previous studies (Illergård et al., 2011), with about 10 % of polar residues in the core membrane regions (see Table 1), and about one-third of these are charged. Over half of the charged residues could form pairs with other charged residues at intervals of *i*, *i*+1; *i*, *i*+3 and *i*, *i*+4. In contrast, the GLOB dataset had a much higher proportion of charged polar residues. The GLOB dataset also contained about 15 times as many charged pairs relative to its size compared to the TM dataset, again indicating that charged residues are more common in globular than in TM helices.

The two datasets, TM and GLOB, were extended with homologous sequences identified by searching with jackhammer against UniProt. All sequences with an E-value lower than 10^-3^ were included. These datasets are named TM-MSA and GLOB-MSA. Using data in the TM-MSA dataset to produce the log odds ratios, we identified periodicity patterns of charged residue pairs that are more common than what would have been expected from the underlying amino acid composition (see Figure 2). We found that polar residues at pairs *i*, *i*+3; *i*, *i*+4 and *i*, *i*+7 are significantly enhanced (Fig. 2). This feature is strengthened when we examine the same plot for the GLOB-MSA dataset, where these patterns were not observed (Figure S2). This was also clear when statistical significance was taken into account, see Figures S3 and S4.

**Figure 2.**
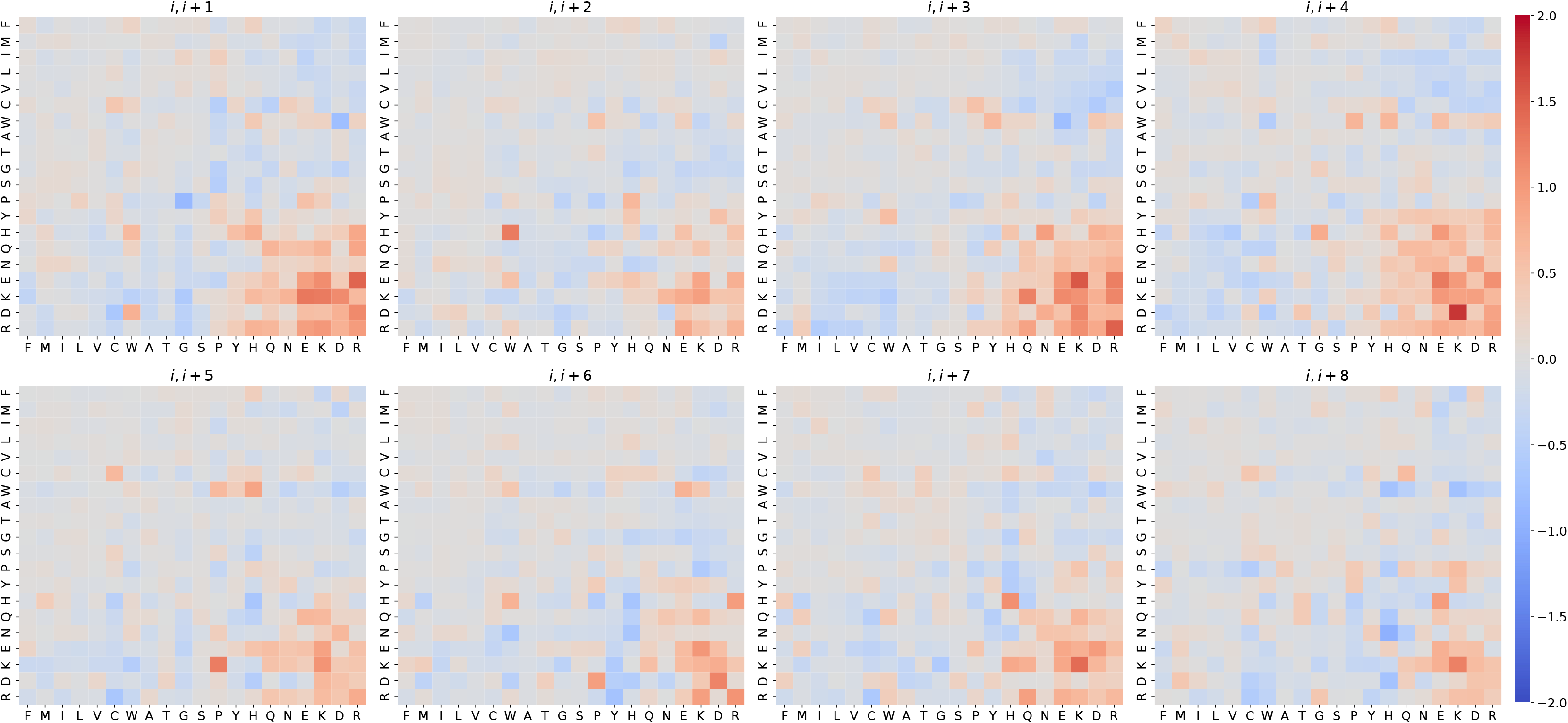
Log odds ratios of each pair of amino acids for ‘*i*, *i*+1’ through ‘*i*, *i*+8’ for the TM-MSA dataset. The rows on the y-axis indicate the first amino acid in the pair and the columns on the x-axis the second. The residues are ordered by hydrophobicity according to the Engelman order (Engelman et al., 1986). See figure S1 for the equivalent of the globular dataset. S2 and S3 show the same plots masked for statistical significance.

Table 2 lists the log odds ratios together with errors and familywise error-corrected *p*-values for all pairs at separation up to seven residues. It is clear that pair residues placed at *i*, *i*+1; *i*, *i*+3 and *i*, *i*+4 positions are most significant, especially in the case of oppositely charged pairs.

**Table 2.** Log odds ratios for all charged pairs, same charged pairs and oppositely charged pairs with calculated errors and multiple hypotheses corrected p-values. It is clear that charged pairs occur more often than predicted, evidenced by the positive log odds ratios in all cases. It is clear that positions +1, +3 and +4 are the most prevalent pairings, with the bolded values highlighting log odds ratio above 0.8. It is also clear that oppositely charged residues which have the potential to form salt bridges are prevalent in all three positions whereas the same charge is mainly prevalent in position 3. Most likely these same charges facing the same face of the helix are involved in functions such as ion transport.

| *Spacing* | *All Log odds* |  |  | *Same charged Log odds* |  |  | *Oppositely charged Log odds* |  |  |
| --- | --- | --- | --- | --- | --- | --- | --- | --- | --- |
|  | *ratio* | *error* | *p-value* | *ratio* | *error* | *p-value* | *ratio* | *error* | *p-value* |
| **+1** | **0.879** | 0.025 | 1.31e<sup>-257</sup> | 0.548 | 0.042 | 4.03e<sup>-41</sup> | **1.107** | 0.032 | 1.65e<sup>-256</sup> |
| **+2** | 0.252 | 0.037 | 2.02e<sup>-08</sup> | 0.185 | 0.054 | 1.88e<sup>-00</sup> | 0.316 | 0.050 | 1.09e<sup>-06</sup> |
| **+3** | **1.073** | 0.026 | <0.00e<sup>-300</sup> | **1.118** | 0.035 | 1.15e<sup>-214</sup> | **1.026** | 0.037 | 5.78e<sup>-165</sup> |
| **+4** | **0.965** | 0.028 | 1.14e<sup>-253</sup> | 0.696 | 0.046 | 2.88e<sup>-49</sup> | **1.177** | 0.036 | 2.74e<sup>-233</sup> |
| **+5** | 0.517 | 0.037 | 4.88e<sup>-42</sup> | 0.390 | 0.055 | 4.15e<sup>-09</sup> | 0.630 | 0.049 | 1.21e<sup>-34</sup> |
| **+6** | 0.540 | 0.038 | 1.66e<sup>-43</sup> | 0.545 | 0.053 | 2.33e<sup>-21</sup> | 0.536 | 0.053 | 2.10e<sup>-20</sup> |
| **+7** | 0.615 | 0.038 | 1.21e<sup>-55</sup> | 0.764 | 0.050 | 7.78e<sup>-50</sup> | 0.439 | 0.058 | 1.97e<sup>-10</sup> |

### Structural analysis of charged residues in transmembrane helices

The abundance of salt bridges also shows a remarkable difference between TM and globular proteins. In the GLOB dataset over 13% of the proteins contains at least one local salt bridge in an alpha helix whereas just over 5% of the membrane proteins contains a local salt bridge, see Table 1. This does conform to our current understanding of soluble versus TM alpha-helices and their different environments, and suggests that salt bridges in TM regions perform a vital functional and/or structural role.

As seen in Figure 2, charged pairs of amino acids are especially prevalent at positions *i*, *i*+1; *i*, *i*+3 and *i*, *i*+4. Oppositely charged residues stand out, especially Glu-Arg at *i*, *i*+1, Glu-Lys at *i*, *i*+3 and Asp-Lys at *i*, *i*+4. Also, several same charged pairings at *i*, *i*+3 and *i*, *i*+6 are more frequent than expected. Other known structural features can also be hinted at, including aromatic ring stacking by His-Trp pair (Samanta et al., 1999) at *i*, *i*+6, and contacting with prosthetic groups by His-His pair at *i*, *i*+7 (Illergård et al., 2011).

Charged pairs placed at *i*, *i*+1; *i*, *i*+3 and *i*, *i*+4 could potentially form salt bridges as they are all on the same relative face of the alpha helix and are close enough in vertical separation on the helix (see Figure 3, top). Although oppositely charged pairs at *i*, *i*+7 are also on the same face of the alpha helix, unless the alpha helix has a bend, both residues are too far separated to form a salt bridge. This was clearly seen in Figure 3 where the TM dataset was used. In both absolute count and log odds ratios (Fig. 3a) it is clear that residues at *i*, *i*+1; *i*, *i*+3 and *i*, *i*+4 are by far the most common and overrepresented pairings. Figure 3a also shows that oppositely and same charged pairs have about the same overrepresentation at *i*, *i*+3, whereas oppositely charged pairs are stronger than same charged pairs at *i*, *i*+1and *i*, *i*+4, both within salt bridge range, and same charged pairs are stronger at positions *i*, *i*+7 and *i*, *i*+8, too far to form salt bridges.

**Figure 3.**
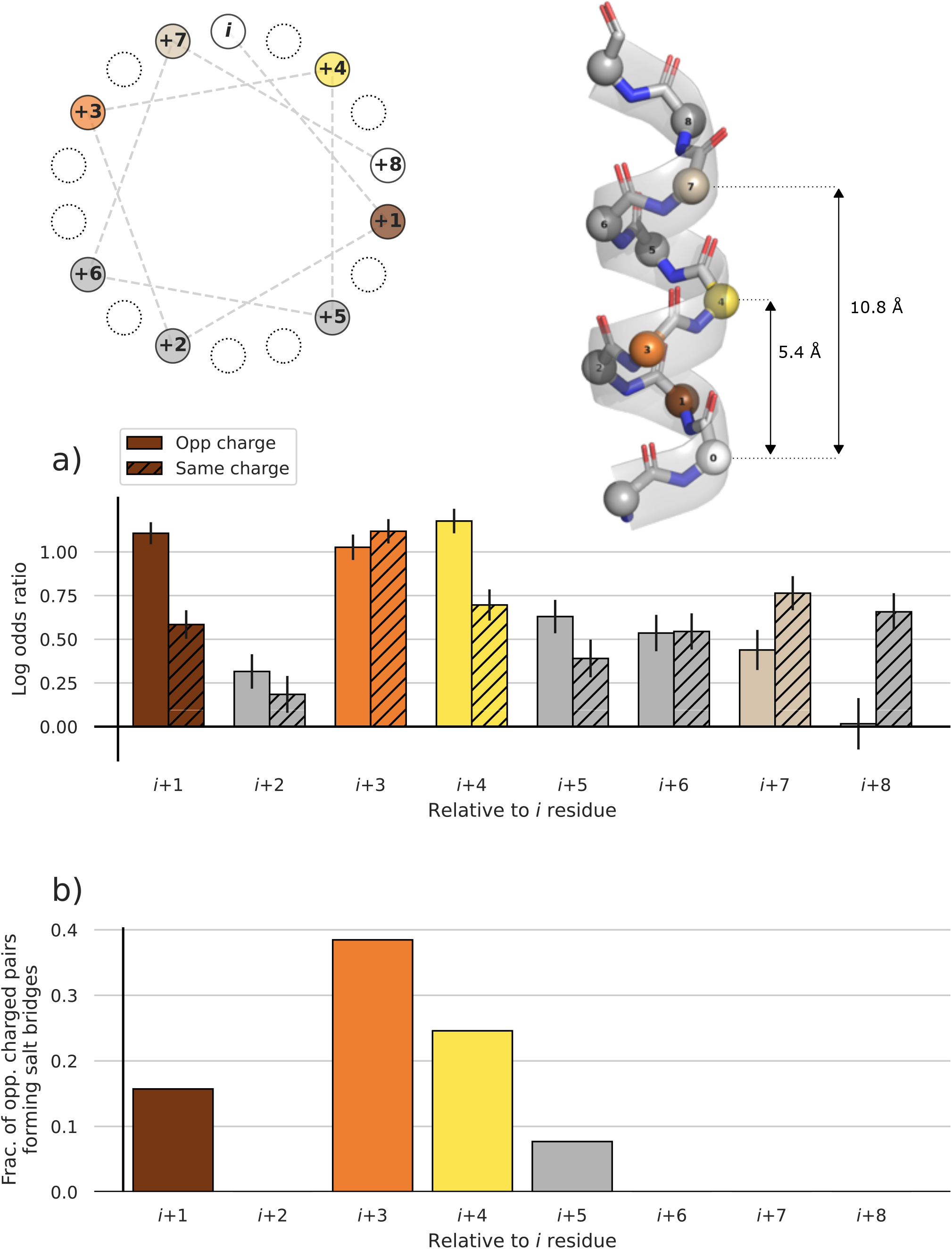
Charge pairs in TM helical sequences and structures. Helical wheel projection and lateral views of an α-helix are shown on top. The initial position *i* and the following 8 residues are numbered. Residues in positions *i*+3 (orange), *i*+4 (yellow), and *i*+7 (light brown) are mainly on the same face of the helix, but *i*+7 is placed too far for a salt bridge interaction. **a)** The log odds ratios of charged pairs for ‘*i*, *i*+1’ through ‘*i*, *i*+8’ in the TM dataset. Plain filled bars refer to oppositely charged pairs and the forward slash are the and same charged pairs, all with error bars. **b)** Fraction of oppositely charged pairs that form local salt bridges. The small bump at *i* +5 are the two proteins 6CC4 (helix A) and 5MG3 (helix E), which both exhibit a bend in the alpha helix due to the presence of glycine residues (see Fig. S6).

When structure-observed salt bridges in the different positions were compared to the oppositely charged pairs a clear image aroused, see Figure 3b. Even though there are more oppositely charged pairs at position *i*, *i*+1 than in positions *i*, *i*+3 and *i, i*+4 with both log odds ratios over 1.0 (Fig. 3a), only about 15% of the oppositely charged pairs at *i*, *i*+1 form salt bridge (Fig. 3b). This is in contrast to *i*, *i*+3 where almost 40 % of the pairs form salt bridges, and just under 25% at *i*, *i*+4 (Fig. 3b).

### Selection of natural salt bridges from membrane protein structures

To further look for candidate proteins containing membrane-spanning helices with salt bridges we started with the redundant (TM-Red) dataset of known structures (see Materials and Methods). Each of the 8,687 proteins in our dataset were scanned for oppositely charged pairs in positions *i, i*+1*; i, i*+3 and *i, i*+4 within any TM segments core region. For each of these, we only kept proteins with at least one TM segment that contains such a pair and where this pair was within salt bridging distance.

To select for potential candidates, we also look at the estimated ΔG_pred_ values and choose helices with a positive value above one, as per our hypothesis, a salt bridge helps TM insertion of sequences for which the hydrophobic force would not be enough to insert. This stricter definition results in a set of 426 candidates of a total of 431 salt bridges with a wide range of estimated ΔG_pred_ penalty values (Figure S5). As shown in Figure S5, most TM segments with a salt bridge exhibit a surprisingly high ΔG_pred_ value above 0 that in normal circumstances are not expected to insert into a membrane. Then, we selected TM7 (helix G) from halorhodopsin protein (PDB ID: 3QBG) with an estimated ΔG_pred_ value above +1.7 kcal/mol, and helix A from calcium ATPase (PDB ID: 1SU4) with a higher estimated ΔG_pred_ value (above +4.1 kcal/mol), as candidates for systematic studies to cover a wide range of insertion penalties. See the github repository (https://github.com/ElofssonLab/salt_bridges) for the full lists.

### Intra-helical salt bridge stabilizes the insertion of helix G from halorhodopsin

Halorhodopsin (hR) from *Natronomonas pharaonis* (3QBG) is a protein made up of seven TM helices (helix A through G) and a retinal chromophore that is bound via a protonated Schiff base to the ɛ-amino group of a lysine (K258) residue located roughly in the middle of helix G (Kanada et al., 2011). *In silico* analysis of 3QBG structure, the anion-free form of the protein, showed a charged pair of amino acids (Asp-Lys), involving the functional K258 and the D254 in the center of helix G (Fig. 4). The position (*i*, *i*+4) and the distance between the anionic carboxylate (RCOO^-^) from the Asp residue and the cationic ammonium (RNH_3_^+^) from the lysine residue, in the crystal structure, was about 3.5 Å, a permissive distance for a salt bridge formation (Fig. 4g), which has been established as being lower than 4 Å (Kumar and Nussinov, 2002). To get insights into this interaction, we designed three mutants that were supposed to perturb the salt bridge interaction by different ways: K258D mutant by placing two charged residues with the same polarity at positions *i*, *i*+4; K258A mutant by replacing one of the charged residues by a non-polar amino acid, and K258Y/Y259K double mutant by placing the charged pair at a non-permissive salt bridge position (*i*, *i*+5; Fig. 4g), while keeping the same amino acid composition (Fig. 4a).

**Figure 4.**
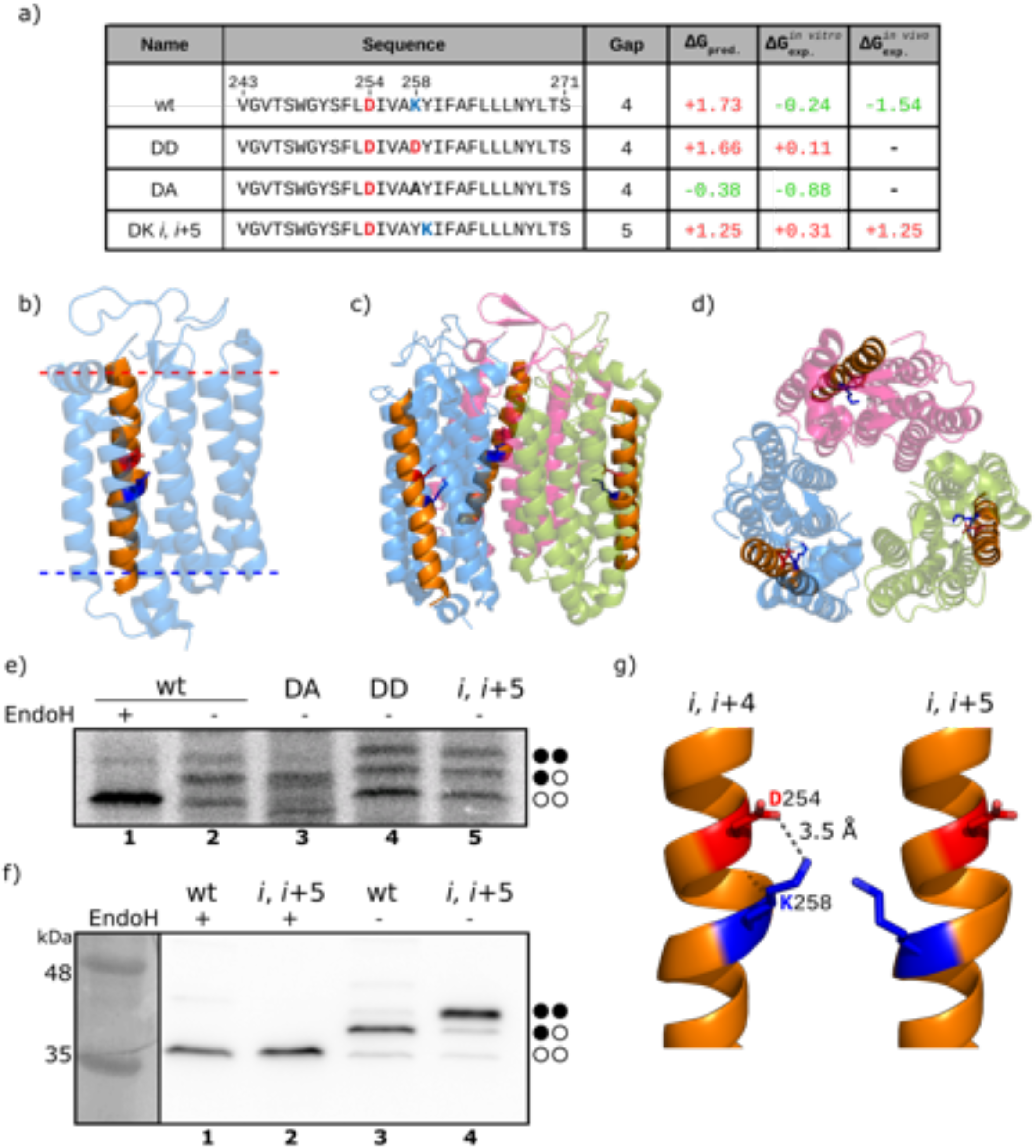
Insertion of halorhodopsin helix G from *Natronomonas pharaonis* (3QBG) into microsomal and cellular membranes. **(a)** Tested sequences from 3QBG including the gap distance, and the predicted (ΔG_pred_) and experimental (*in vitro* ΔG_exp_*^in vitro^* and in vivo ΔG_exp_*^in vivo^*, respectively) ΔG values in kcal/mol are shown. Amino acids with a positive or negative charge are highlighted in blue (K) and red (D), respectively. Green numbers indicate negative ΔG (insertion) values, while red numbers denote ΔG values above 0 (translocation). **(b)** Frontal view of 3QBG monomer structure. The helix G is highlighted in orange with the D254 and K258 shown in sticks colored red and blue respectively. The membrane position is indicated by a red (outer) and blue (inner) discontinuous line, according to OPM dataset [22]. Lateral **(c)** and upper **(d)** views of the 3QBG trimeric structure. The helix G is highlighted in orange with the D and K shown in sticks colored red and blue, respectively. The different monomers are shown in transparent blue, pink and green. Representative examples (*n*=3) of *in vitro* protein translations in the presence of ER-derived microsomes **(e)** and Western blots (*n*=3) of *in vivo* protein translations in HEK-293T cells **(f)** in the presence (+) or absence (-) of Endoglycosidase H (EndoH), a glycan-removing enzyme. The absence of glycosylation of G1 and G2 acceptor sites is indicated by two white dots, single glycosylation by one white and one black dot, and double glycosylation by two black dots. **(g)** Zoom view centered on the salt bridge between D254 and K257 at *i*, *i*+4 (left) and *i*, *i*+5 (right) gaps. D and K residues are shown in sticks colored red and blue, respectively, while the dashed line indicates the RCOO^-^ to RNH_3_^+^ distance.

Halorhodopsin is a trimeric protein in which the helix G is neither exposed at the monomer-monomer interface nor oriented to the inner part of the trimeric structure (Fig. 4, panels b-d). In fact, the charged pair found in helix G is oriented toward the core of the ‘globular’ structure in each monomer. Then, the insertion of helix G was studied using the Lep-based assay. When constructs harboring helix G wild type sequence were translated *in vitro* in the presence of RMs singly-glycosylated (reporting insertion) forms were found (Fig. 4e, lane 2), despite its positive (suggesting non-insertion) ΔG_pred_ value (Fig. 4a). The nature of the higher molecular weight polypeptide species was analysed by endoglycosidase H (EndoH) treatment, a highly specific enzyme that cleaves N-linked oligosaccharides. Treatment with EndoH of the samples eliminated higher molecular mass bands (Fig. 4E, lane 1), confirming the sugar source of the retarded electrophoretic mobility bands and suggesting helix G insertion into the microsomal membrane. However, locating the Asp-Lys pair at *i*, *i*+5, which is non-compatible with salt bridge interaction (Fig. 4g), strongly reduced the experimental insertion efficiency (Fig. 4e, lane 5). Interestingly, replacing the positively charged lysine residue by a negatively charged aspartic acid residue (K258D) rendered similarly low levels of insertion efficiency (Fig. 4e, lane 4). As expected, replacement of the ionizable lysine residue by the aliphatic alanine increased the insertion efficiency (Fig. 4e, lane 3). In this later mutant, the most likely cause for the increased insertion is the absence of the positively charged lysine amino side chain and by the presence of the methyl side chain group of the mutant alanine residue. The results of the Lep-based glycosylation assay indicated that wild type (wt) and DA sequences (two stabilized charges or only one charge, respectively) are inserted properly into the microsomal membrane (ΔG_exp_ values −0.24 and −0.88 kcal/mol, respectively), but when the salt bridge is disrupted, either by having two charged amino acids with the same polarity (DD) or by placing oppositely charged residues at a non-permissive distance (*i*, *i*+5) in the center of the helix, the translocation of the segment increases substantially. It should be mentioned that the K258Y/Y259K double mutant has the same amino acid composition than the original helix G, but insertion efficiency is remarkably decreased (ΔG_exp_ = +0.31 kcal/mol). Together these results show that the interaction (salt bridge) between Asp and Lys residues in the center of the helix G from 3QBG is essential for its proper insertion into the microsomal membrane. The salt bridge contributes approximately ∼0,5 kcal/mol to the apparent experimental free energy of microsomal membrane insertion, as this is the difference found between the ΔG_exp_ values for the wt and *i*, *i*+5 mutant.

Next, to ensure that the *in vitro* results are relevant to the *in vivo* situation, wt and *i, i*+5 constructs were also expressed *in vivo* in HEK-293T cells. To this end, a *c*-myc tag was engineered at the C-terminus of the Lep chimera to allow immune-detection of our constructs in cell extracts. As shown in Fig. 4f, transfected cells with the chimera containing helix G wt sequence rendered singly glycosylated molecules, indicating *in vivo* membrane insertion. In contrast, cells transfected with the construct harboring *i*, *i*+5 sequence rendered almost exclusively doubly glycosylated forms, as proved by EndoH treatment (Fig. 4f, lane 2), suggesting membrane translocation. These results emphasized the relevance of salt bridge interactions in translocon-mediated TM insertion, especially in the *in vivo* (cellular) environment.

### Salt bridge contribution to the insertion of a heavily charged helix

In order to challenge salt bridge interactions in a more hydrophilic TM helix (Figure S5), we focus on helix A from the sarcoplasmic/ER calcium ATPase 1 (Toyoshima et al., 2000). Calcium ATPase (PDB ID: 1SU4, *Oryctolagus cuniculus*) is a member of the P-type ATPases that transport ions across the membrane against a concentration gradient involving 10 α-helices (helix A through J) in the membrane-embedded region (Toyoshima et al., 2000). *In silico* analysis of 1SU4 structure, the crystal structure of the calcium ATPase with two bound calcium ions, showed Asp-Arg (DR) pair (D59 and R63) in the center of the helix A. This helix extends beyond the membrane (Fig. 5a, from Leu49 to Phe78) and shows some particularities. On the one hand, in the structure the membrane-embedded stretch (the N-terminal region of helix A) encompasses from Leu49 to Ala69 residues, and includes several charged amino acids, probably involved in the Ca^2+^ transport across the membrane (Glu51, Glu55, Glu58, Asp59 and Arg63). Therefore, the ΔG_pred_ value for this segment (L49-A69) is remarkably higher (and positive, +4.12 kcal/mol) than expected for a TM helix (Fig. 5b). On the other hand, the C-terminal region of this helix contains a high prevalence of non-polar amino acids that is more compatible with the hydrophobicity of the membrane core. Nevertheless, the presence of the functional amino acids (e.g. Glu58) in the membrane-embedded region reinforce the idea that the more hydrophilic N-terminal region must ultimately be embedded within the membrane (Toyoshima et al., 2000). It has been previously shown that the position in the membrane of TM helices in protein folded structures does not always correspond to the thermodynamically favored positions in the membrane of the isolated helices (Kauko et al., 2010). Instead, after translocon-mediated insertion of the more hydrophobic region, repositioning of TM helices relative to the lipid bilayer provides a convenient way for non-hydrophobic polypeptide segments to become buried within the membrane. Then, the nature of helix A suggested the possibility that initial insertion of the hydrophobic region can be followed by subsequent repositioning of the charged region into the membrane hydrophobic core, which will be the final segment embedded in the lipid bilayer (Fig. 5a). Interestingly, the ΔG Predictor server (https://dgpred.cbr.su.se/index.php?p=home) selected the adjacent (C-terminus, L60-F78) hydrophobic region as TM (Fig. 5b), instead of the charged region found at the high-resolution structure (Uniprot code: P04191).

**Figure 5.**
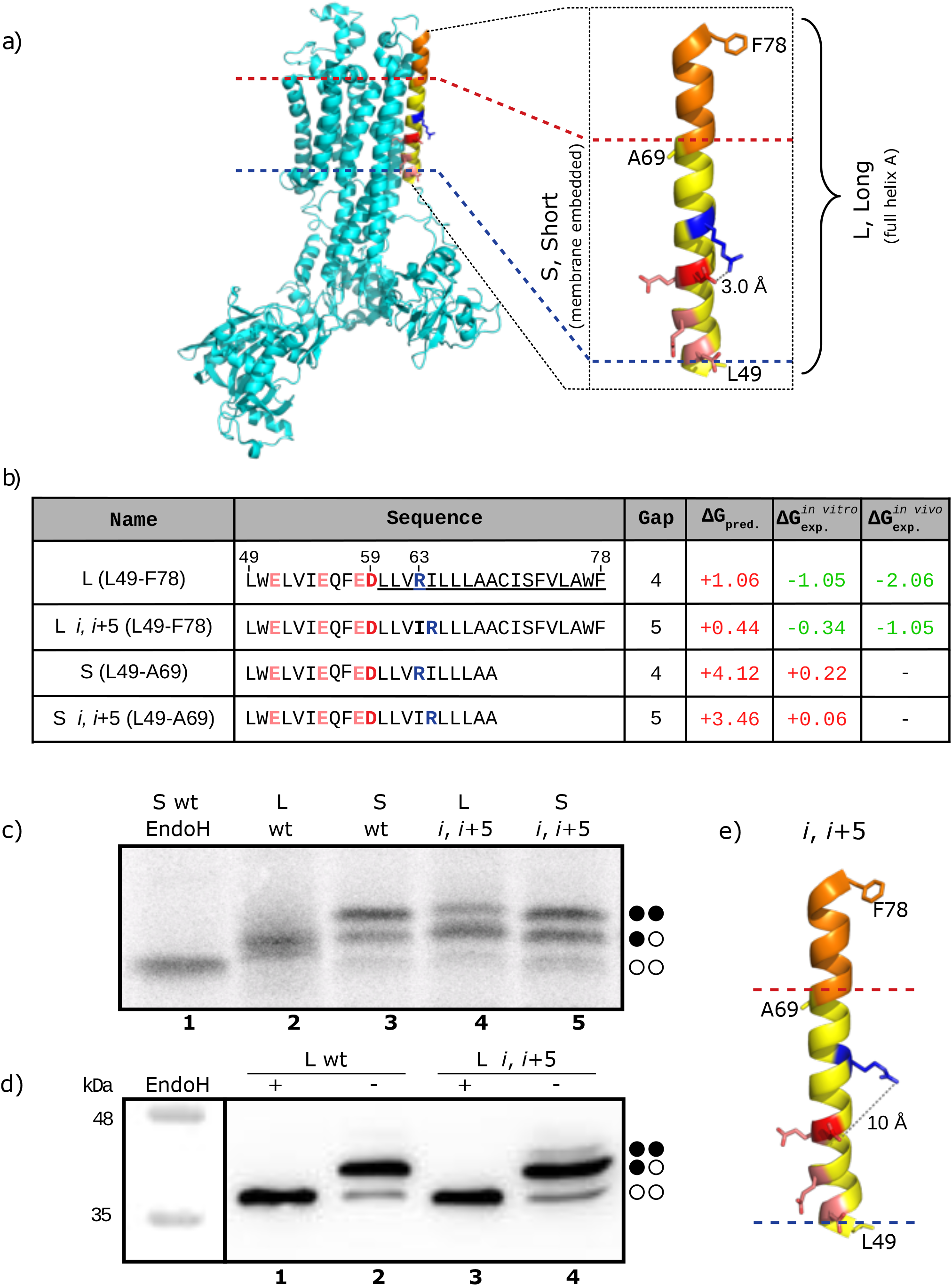
Insertion of Calcium ATPase (1SU4) helix A into microsomal and cellular membranes. **(a)** Lateral view of 1SU4 structure. Zoom view of the A helix (right panel). The membrane-embedded region of helix A is highlighted in yellow. Charged amino acids are shown as sticks in blue (R), red (D) and pink (E), respectively. L49, A69 and F78 are also shown as sticks to define helix’s subdomains. The membrane location is indicated by red (outer) and blue (inner) discontinuous lines according to OPM dataset [22] and the distance between the R and D charges is indicated in Å. **(b)** Helix A-derived sequences from 1SU4 including the gap between charged residues, and the predicted (ΔG_pred_) and experimental (*in vitro* ΔG_exp_*^in vitro^* and *in vivo* ΔG_exp_*^in vivo^*, respectively) ΔG values in kcal/mol are shown. Amino acids with a positive charge are highlighted in blue (K) while negatively charged are marked in red (D) and pink (E). The residues predicted as TM by the ΔG Prediction server are underlined. Green numbers indicate negative ΔG (insertion) values while red numbers denote ΔG values above 0 (translocation). Representative examples (n=3) of *in vitro* protein translations in the presence of ER-derived microsomes **(c)** and Western blots (n=3) of *in vivo* protein translation in HEK-293T cells **(d)** in the presence (+) or absence (-) of Endoglycosidase H (EndoH), a glycan-removing enzyme. The absence of glycosylation of G1 and G2 acceptor sites is indicated by two white dots, single glycosylation by one white and one black dot, and double glycosylation by two black dots. **(e)** 1SU4 helix A *i*, *i*+5 mutant. The membrane-embedded region of helix A is highlighted in yellow. Charged residues are shown as sticks in blue (R), red (D) and pink (E). L49, A69 and F78 are also shown as sticks to define helix’s subdomains. The membrane is indicated by red (outer) and blue (inner) discontinuous lines as in (a), and the dashed line indicates the RCOO^-^ and RC(NH_2_)_2_^+^ distance.

Focusing on the potential salt bridge residues (D59 and R63 pair), the distance between the anionic carboxylate (RCOO^-^) from the D59 and the cationic guanidinium (RC(NH_2_)_2_^+^) from the R63 was about 3.0 Å in the crystal structure (Figure 5a), clearly within the permissive range for salt bridge formation. To investigate the contribution of this potential salt bridge interaction in the translocon-mediated membrane insertion of this region, we worked with two different scaffold sequences: the full helix A involving the residues 49-78 (Long, L); and a shorter membrane-embedded version including residues 49-69 (Short, S) as found in the solved structure. We also challenged the D59-R63 charge pair interaction in both sequences by increasing the separation between the ionizable residues from the native *i*, *i*+4 to non-permissive *i*, *i*+5, while maintaining amino acid composition (R63I/I64R double mutant). *In vitro* transcription/translation of these sequences in the presence of microsomes rendered singly glycosylated molecules for the construct containing full-length helix A (Fig. 5c, lane 2). In contrast, when only the membrane-embedded sequence was included, the Lep chimera was mainly doubly glycosylated (Fig. 5c, lane 3), suggesting that in the full-length protein helix A inserts initially through the more hydrophobic L60-F78 region and then, after protein rearrangements, repositions the more hydrophilic L49-A69 region at the membrane core, as found in the solved structure. Accordingly, the translocon inefficiently inserted the isolated membrane-embedded (L49-A69) region (Fig. 5c, lane 3), properly inserting helix A only when the full helical sequence is present (Fig. 5c, lane 2). When the charge paired residues were placed at a non-permissive distance in terms of salt bridge interaction (*i*, *i*+5; Fig. 5e) the insertion efficiency was reduced (Fig. 5c, lane 4), with a ΔG_app_ decrease (absolute values) of ∼0,7 kcal/mol relative to the wild type sequence (Fig. 5b). As expected, this effect was not observed when the same mutations were grafted on the membrane-embedded (S) sequence (Fig. 5c, lane 5).

Next, we analysed the salt bridge interaction in HEK-293T cells to study the translocon performance *in vivo*. When cell cultures were transfected with a Lep-derived chimera containing the helix A wild type sequence only singly glycosylated molecules were observed (Fig. 5d, lanes 1 and 2). However, a construct harboring double mutant (R63I/I64R; *i*, *i*+5) sequence showed doubly glycosylated molecules to a measurable extent (Fig. 5d, lanes 3 and 4), indicating a lower insertion efficiency that can be attributed to the altered salt bridge interaction.

### Effect of salt bridge formation in the absence of previous TM regions

To investigate the effect of salt bridge formation in translocon-mediated membrane insertion in the absence of precedent TM regions we used a different glycosylation-based reporter system in which the TM sequences of interest (bR helix G and ATPase helix A) were connected to the well-folded constant domain of the antibody λ light chain, C_L_ (Feige and Hendershot, 2013). The C-terminal of the TM sequence carries an Asn-Val-Thr glycosylation site (G2), and we engineered an extra glycosylation site (G1) within the C_L_ sequence (Figure 6a, top). As in the case of the Lep system, G1 will always be glycosylated due to the native translocation of the λ light chain, but G2 will be glycosylated only upon translocation of the analyzed (tested) sequence across the ER membrane (Figure 6a, bottom). When constructs harboring either hR helix G or ATPase helix A wild type sequences were transfected into Hek293T cells, singly glycosylated (reporting insertion) forms were predominantly found (Fig. 6b, lane 1 and Fig. 6c, lane 2, respectively), discarding any potential contribution to membrane insertion of precedent TM segments. On the contrary, sequences containing non-permissive (*i*, *i*+5) salt-bridge forming pairs were more efficiently doubly glycosylated (Figures 6b, lane 3 and 6c, lane 4, respectively). Thus, similar to what was observed using the Lep system (Figures 4f and 5d), the presence of chair pairs at permissive salt bridge formation distances strongly promotes helix integration into the ER membrane.

**Figure 6.**
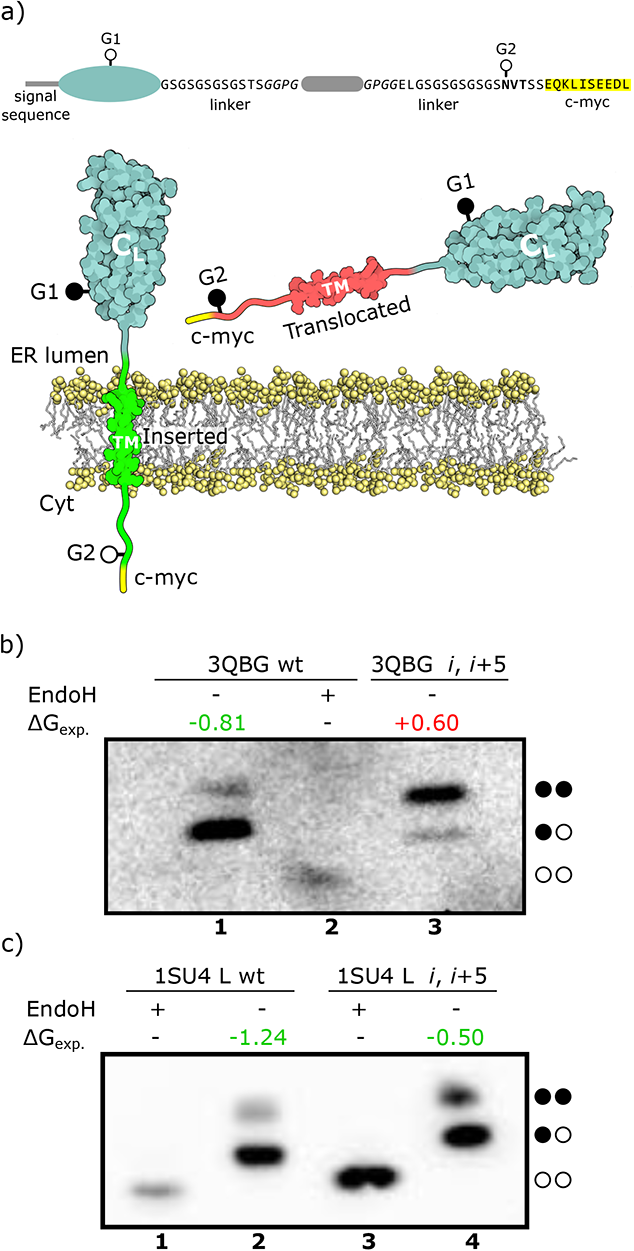
Salt bridge effect on the insertion in the absence of preceding TM segments. **(a)** (Top) Schematic of the C_L_TM construct used. It is composed of the domain of the antibody λ light chain (C_L_) containing a glycosylation site (G1) connected by flexible linkers to the sequence of analysis followed by a C-terminal glycosylation site (G2) and a *c*-myc tag. (Bottom) Scheme depicting the main features of the C_L_TM insertion assay. Black dots represent glycosylated sites while white dots represent non-glycosylated sites. **(b)** Representative halorhodopsin (3QBG) helix G western blot (*n*=3) of *in vivo* protein translations in HEK-293T cells in the presence (+) or absence (-) of Endoglycosidase H (EndoH), a glycan-removing enzyme. **(c)** Representative Ca^2+^ ATPase (1SU4) helix A western blot (*n*=3) of *in vivo* protein translations in HEK-293T cells in the presence (+) or absence (-) of EndoH. Non-glycosylated proteins are indicated by two white dots, singly-glycosylated proteins are indicated by one white and one black dot, and doubly-glycosylated proteins are indicated by two black dots. Experimental ΔG values (kcal/mol) are shown above each sample (*n*=3).

## DISCUSSION

Charged residues found in alpha-helices can both give a stabilizing effect (Armstrong and Baldwin, 1993) as well as being important for interaction and function such as in zinc finger motifs (Lin and Lin, 2018). Charged residues have also been found to be more common in globular (soluble) helices compared to TM helices (Kauko et al., 2008). This can be explained by the fact that globular helices reside in a more hydrophilic environment compared to the hydrophobic environment of the lipid bilayer. An intermediate between these is the existence of amphipathic alpha-helix in contact with the surface of a bilayer, where one face contains mainly polar residues facing an aqueous environment and the opposite face with mostly nonpolar residues facing a hydrophobic environment (Giménez-Andrés et al., 2018).

Whereas pairs of charged residues of the same charge facing the inside pores in TM regions fill an essential functional role, pairs of oppositely charged residues can form salt bridges that can stabilize the helix and play an important role in function such as for transportation (Walther and Ulrich, 2014). Salt bridges might also be important for hiding the charges during translocon-mediated TM helix insertion, as the charged residues are hidden from the hydrophobic environment. Previous work has shown that charged and polar residues are conserved within TM segments (Illergård et al., 2011), indicating they are crucial for function and/or stability.

Asp-Lys pairs at position *i*, *i*+4 and Glu-Lys pairs at position *i*, *i*+3 are the most prevalent as seen previously in Figure 2. They are both among the most prevalent oppositely charged pairs and the charged pairs that form the highest number of salt bridges in membrane protein structures. This is in stark contrast to Glu-Arg pair at position *i*, *i*+1 that although as frequent in pairs as Asp-Lys and Glu-Lys at positions *i*, *i*+4 and *i*, *i*+3 respectively, only form salt bridges in one-fourth of the cases as found in Fig. 3b.

An interesting observation is that positively charged Arg-Arg pairs at position *i*, *i*+3 is numerically the most common pair at this position. Positively charged pairs at *i*, *i*+6 are also more prevalent than expected and although Arg-Arg pairs numerically only show up half as often at *i*, *i*+6 as in *i*, *i*+3, 80% of the Arg-Arg pairs at position *i*, *i*+6 also contain an arginine at *i*+3, as found in helices from voltage-gated ion channels that contain three or more periodically aligned Arg residues with two intervening hydrophobic residues (Okamura et al., 2015).

The fraction of salt bridges at positions *i*, *i*+5 (Fig. 3b) is an anomaly due to bend alpha-helices, as found in the helix E from bacterial translocon (PDB ID: 5MG3) and in the helix A from a lipid flippase (PDB ID: 6CC4) due in both cases to the presence of a glycine residue (Figure S6). Without the observed helix bending, these salt bridges would not be formed.

By analyzing two native helices containing intra-helical salt bridges we now find that the free energy of insertion (ΔG_app_) is significantly reduced if both oppositely charged residues are spaced at a permissive distance. These results indicate that intra-helix salt bridge can form during translocon-assisted insertion or even earlier, since in contrast to globular (soluble) helices, TM helices can be compacted inside the ribosome exit tunnel (Bañó-Polo et al., 2018). The maximal reduction in ΔG_app_ seen with Asp-Lys and Asp-Arg pairs in both hR and Ca^2+^ ATPase helices is 0.5-0.7 kcal/mol, which is in good agreement with the 0-1 kcal/mol estimated for these two pairs from thermodynamic peptide partition into octanol experiments (Jayasinghe et al., 2001). As found in the case of hR helix G (Figures 4f and 6b), this reduction might be even higher in the cell context, since some auxiliary components of the membrane insertion machinery (Chitwood and Hegde, 2020; Shurtleff et al., 2018; Tamborero et al., 2011) can be in suboptimal conditions in the microsomal vesicles.

As mentioned above, in the case of hR helix G the lysine residue involved in the salt bridge (K258) is bound to the retinal chromophore via a protonated Schiff base as found in the crystal structure of a close homologue (Kolbe et al., 2000). Then, the lysine residue plays a fundamental role for protein function but at the same time introduces a penalty for membrane insertion. Interestingly, our data suggest that helix G translocon-mediated insertion efficiency could be increased by salt bridge formation between K258 and D254, and once in the membrane, retinol binding to the apoprotein could occur by covalently binding the retinal as a protonated Schiff base to K258 and perturbing D254 salt bridge interaction.

Beyond the conceptual issues involving the membrane insertion process, we note that the availability of quantitative experimental data on the contribution of salt bridge interactions to the free energy of insertion (ΔG_app_) will make it possible to fine-tune current membrane protein topology-prediction methods based on free energy calculations. Although today’s state of the art topology prediction tools uses amphiphatic biologically derived scales, they do not take these types of salt bridge interaction into account. Current algorithms tend to overestimate the free energy of insertion due to the penalty acquainted by charged residues in the TM region. However, distinguishing between charged residues of the same or opposite polarity, i.e., incorporating the effect of potential salt bridges in the reduction of ΔG_app_ during membrane integration should help to make prediction tools even more accurate.

## MATERIALS AND METHODS

### Enzymes and chemicals

TNT T7 Quick for PCR DNA was from Promega (Madison, WI, USA). Dog pancreas ER column washed rough microsomes were from tRNA Probes (College Station, TX, USA). EasyTag™ EXPRESS^35^S Protein Labeling Mix, [^35^S]-L-methionine and [^35^S]-L-cysteine, for *in vitro* labeling was purchased from Perkin Elmer (Waltham, MA, USA). Restriction enzymes were from New England Biolabs (Massachusetts, USA) and endoglycosidase H was from Roche Molecular Biochemicals (Basel, Switzerland). PCR and plasmid purification kits were from Thermo Fisher Scientific (Ulm, Germany). All oligonucleotides were purchased from Macrogen (Seoul, South Korea).

### DNA Manipulation

The sequences of interest were introduced into the modified Lep sequence from the pGEM1 plasmid (Hessa et al., 2005) between the *Spe*I and *Kpn*I sites using two double-stranded oligonucleotides with overlapping overhangs at the ends. The complementary oligonucleotides pairs were first annealed at 85 °C for 10 min followed by gradual cooling to 30 °C and ligated into the vector (a kind gift from G. von Heijne’s lab). Mutations were obtained by site-directed mutagenesis using the QuikChange kit (Stratagene, La Jolla, California). Lep system including the sequences of interest in the P2 region were subcloned into *Kpn*I linearized pCAGGS in-house version using In-Fusion HD cloning Kit (Takara) according to the manufacturer’s instructions. An engineered glycosylation site (Q36N) was added to the C_L_-TM plasmid (a kind gift from L. Hendershot’s lab), in which the sequences from hR helix G and ATPase helix A were introduced flanked by ‘insulating’ Gly-Pro tetrapeptides. A *c*-myc tag (EQKLISEEDL) at the C-terminus of the Lep-and C_L_-derived sequences was added by PCR before cloning. For *in vitro* assays, DNA was amplified by PCR adding the T7 promoter during the process. All sequences were confirmed by sequencing the plasmid DNA at Macrogen Company (Seoul, South Korea).

### Translocon-mediated insertion into microsomal membranes

Lep constructs in pGEM with L4/A15, L5/A14, 3QBG and 1SU4 segments and its variations were transcribed and translated using the TNT T7 Quick Coupled System (#L1170, Promega). Each reaction containing 1 μL of PCR product, 0.5 of EasyTag^TM^ EXPRESS 35S Protein Labeling Mix (Perkin Elmer) (5.5 μCi) and 0.3 μL of microsomes (tRNA Probes) was incubated at 30°C for 90 min. Endo H treatment was done following the manufacturer’s instructions. Samples were analysed by SDS-PAGE (12-14% polyacrylamide). The bands were quantified using a Fuji FLA-3000 phosphoimager and the Image Reader 8.1j software. Free energy was calculated using: ΔG_app_=-RT lnK_app_, where K_app_=f_2g_/f_1g_ being f_1g_ and f_2g_ the fraction of singly glycosylated and double glycosylated protein, respectively.

### Free apparent insertion energy, ΔG

The free insertion energy of a TM region, ΔG, is calculated as per the experimentally defined Biological hydrophobicity scale (Hessa et al., 2005). This scoring is amphiphilic with hydrophobic residues contributing a lower (negative) ΔG while hydrophilic contributes a higher (positive) ΔG. The total ΔG of a region is the sum of individual position specific scores. This scoring can give an indication of how favorable the amino acid composition of a TM region is to be inserted in a lipid bilayer membrane. To note is that hydrophobicity alone is not the only driving force and that the positive inside rule (Heijne, 1989; Lerch-Bader et al., 2008) and help from proceeding TM regions (Bañó-Polo et al., 2013; Hedin et al., 2010) can assist insertion in polytopic TM proteins especially.

### Core segment definition

We define core segments as a TM region minus the first and last 5 residues. This is to ignore the interface regions which are known to contain polar residues.

### Salt bridge definition

Salt bridges are defined as per (Kumar et al., 2000), where a salt bridge is defined if a side chain carbonyl oxygen atom in Asp-Glu is within 4.0 Å from the nitrogen atom in Arg-Lys. This conforms to other works (Bosshard et al., 2004; Donald et al., 2011) with the definition that the atoms are within hydrogen bond distance. We also define local salt bridges as being bridges that are separated by at most 7 residues in the sequence. This is to separate long salt bridge interactions, which can occur between spatially close residues that are separated in sequence, such as coiled coils where salt bridges can be between separate alpha-helices.

### Transmembrane helices dataset (TM dataset)

The full pipeline is available as a Makefile together with supporting scripts in the github repository. The full PDBTM database (Kozma et al., 2013) was downloaded together with their list of non-redundant protein pdb ids. This list is used to generate both sequence and topology of the proteins by extracting both from the PDBTM-xml. For each protein in the non-redundant list, the membrane regions are extracted as per the PDBTM database annotation. All non-membrane regions are annotated ‘i’ for convenience. To support future analysis, membrane regions longer than 10 were run through DSSP and annotated with ‘M’ if all residues in the core segment were defined as alpha-helix (‘H’), otherwise, the full membrane region is annotated ‘m’. Observe that this creates fasta-like 3line files, that only contain topological annotation with ambiguous TM regions annotated as ‘m’ instead of the normal ‘M’. These proteins are then cluster using cd-hit (Fu et al., 2012; Li and Godzik, 2006) at 40% identity using the parameter -c 0.4 -n 2 -T 0 -M 0 -d 0.

During the extraction of TM regions, the corresponding structure file from RCSB was used to calculate all salt bridges within the current protein and any salt bridge that has at least one residue within any TM region was saved. Additionally, all salt bridges whose both residues were within the same segment and within 7 residues of each other were annotated as local as per the salt bridge definition above.

### Extraction of charged residues

From all annotated TM regions of length 17 or longer (Baeza-Delgado et al., 2012), the core segment was extracted. All these core regions were then scanned and when a charged residue was encountered, we recorded any other charged residue from 7 residues before the current one to 7 residues after. This results in charged residues that can contain a charged pairing partner outside of the TM segment and will therefore differ slightly from charged pairs which are defined next.

### Extraction of charged pairs

From all annotated TM regions of length 17 or longer (Baeza-Delgado et al., 2012), the core segment was extracted. All these core regions were then scanned and when a charged residue was encountered, we look at up to 7 residues in front of it or to the end of that core region, whichever came first. All occurrences of charged pairs were recorded, resulting in charged pairs where both residues were fully within the core segment of a TM helix.

### MSA-dataset extension (TM-MSA)

Using the TM dataset, we extended it by creating an MSA alignment of each protein using jackhmmer (Eddy, 2011) against Uniref90 with one iteration and an E-value cut off of 10^-3^ with the following parameters:

-N 1 -E 1e-3 --incE 1e-3 --cpu 14

From each alignment, we then sampled up to 200 hits, including the initial seed sequence. If there were fewer than 200 hits, we used them all. We then used the original topology for each alignment to extract all TM regions, only to include parts where the sequence covers the full TM region and where the sequence did not contain any insertions or deletions.

### Dataset of helices from globular proteins (GLOB and GLOB-MSA)

To create a reference dataset of globular α-helical proteins we extracted all globular all-alpha protein domains from SCOP. As SCOP classifies domains of proteins resulting in that one domain of a protein can be annotated as globular whereas another domain as TM (see 1PPJ chain D as an example) we reduced the SCOP list against the redundant list of all PDBTM (Kozma et al., 2013) chains to clear any overlap. This results in 4,500 proteins in total.

Topological files with sequence and membrane topology are created with the help of the RCSB secondary structure file and only membrane segments whose core region (*i.e.*, central 15 residues) is annotated as pure (canonical) α-helices were retained, *i.e.*, those TM segments containing any residues within the core annotated as loops or other types of secondary structures were removed. This file was further homology reduced and alignments prepared in the same manner and using the same parameters as the TM dataset described above.

The GLOB-MSA dataset was created in the same way as the TM-MSA dataset using jackhmmer to extend the sequences to alignments and then to extract helix sequences.

### Redundant (TM-Red) dataset

The full PDBTM database was used as in the preparation of the TM dataset. We skip the clustering step and instead use all redundant proteins to generate their respective topology files. We added in the constraint that each selected TM region must fully contain at least one potential salt bridge. This means a local salt bridge where both residues are within the core segment. This dataset was only used to find potential candidates for further *in vitro* and *in vivo* experiments. See section of natural salt bridges above.

### Calculation of log odds ratio

The log odds ratios for each amino acid pair for steps 1 through 10 are calculated as follows:

*logOddsRatio = log((A/B)/(C/D))*

Where for two amino acids p_1_ and p_2_:

*A* = number of pairs of p_1_ to p_2_

*B* = number of total pairs

*C* = number of p_1_ times number of p_2_

*D* = number of total pairs squared

The standard error, SE, and z-value is calculated as follows allowing for a two-sided test:

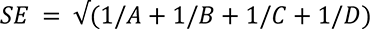

*z = abs(logOddsValue/SE)*

The survival function, sf, from the scipy python packages is used to calculate the p-values. To correct for multiple hypothesis, the Bonferroni Correction is used based on the number of hypothesis, 20 * 20 * 10, number of amino acids square times the number of steps.

### Expression in mammalian cells

Lep or C_L_-derived constructs containing 3QBG or 1SU4 segments and its variations were tagged with *c*-myc epitope at their Ct (EQKLISEEDL) and inserted in the appropriate plasmids. Once the sequence was verified, plasmids were transfected into HEK293-T cells using Lipofectamine 2000 (Life Technologies) according to the manufacturer’s protocol. Approximately 24 h post-transfection cells were harvested and washed with PBS buffer. After a short centrifugation (1000 rpm for 5 min on a table-top centrifuge) cells were lysed by adding 100 μL of lysis buffer (30 mM Tris-HCl, 150 mM NaCl, 0.5% Nonidet P-40) were sonicated in an ice bath in a bioruptor (Diagenode) during 10min and centrifuged. After protein quantification, equal amounts of protein were submitted to Endo H treatment or mock-treated followed by SDS-PAGE analysis and transferred into a PVDF transfer membrane (ThermoFisher Scientific) as previously described (Duart et al., 2020). Protein glycosylation status was analysed by Western Blot using an anti-*c*-myc antibody (Sigma), anti-rabbit IgG-peroxidase conjugated (Sigma), and with ECL developing reagent (GE Healthcare). Chemiluminescence was visualized using an ImageQuantTM LAS 4000mini Biomolecular Imager (GE Healthcare).

## Acknowledgments

We thank Pilar Selvi and Beatriz Iborra for excellent technical and administrative assistance, respectively. The C_L_-TM plasmid was a kind gift from Prof. Linda M. Hendershot (St. Jude Children’s Research Hospital). This work was supported by grants PID2020-119111GB-100 from the Spanish Ministry of Science and Innovation and PROMETEU/2019/065 from Generalitat Valenciana (to I.M.), and by grant from the Swedish Research Council (VR-NT 2016-03798 to A.E.). G.D. was recipient of a predoctoral contract (FPU18/05771) from the Spanish Ministry of Education.

---

**Figure S1.**
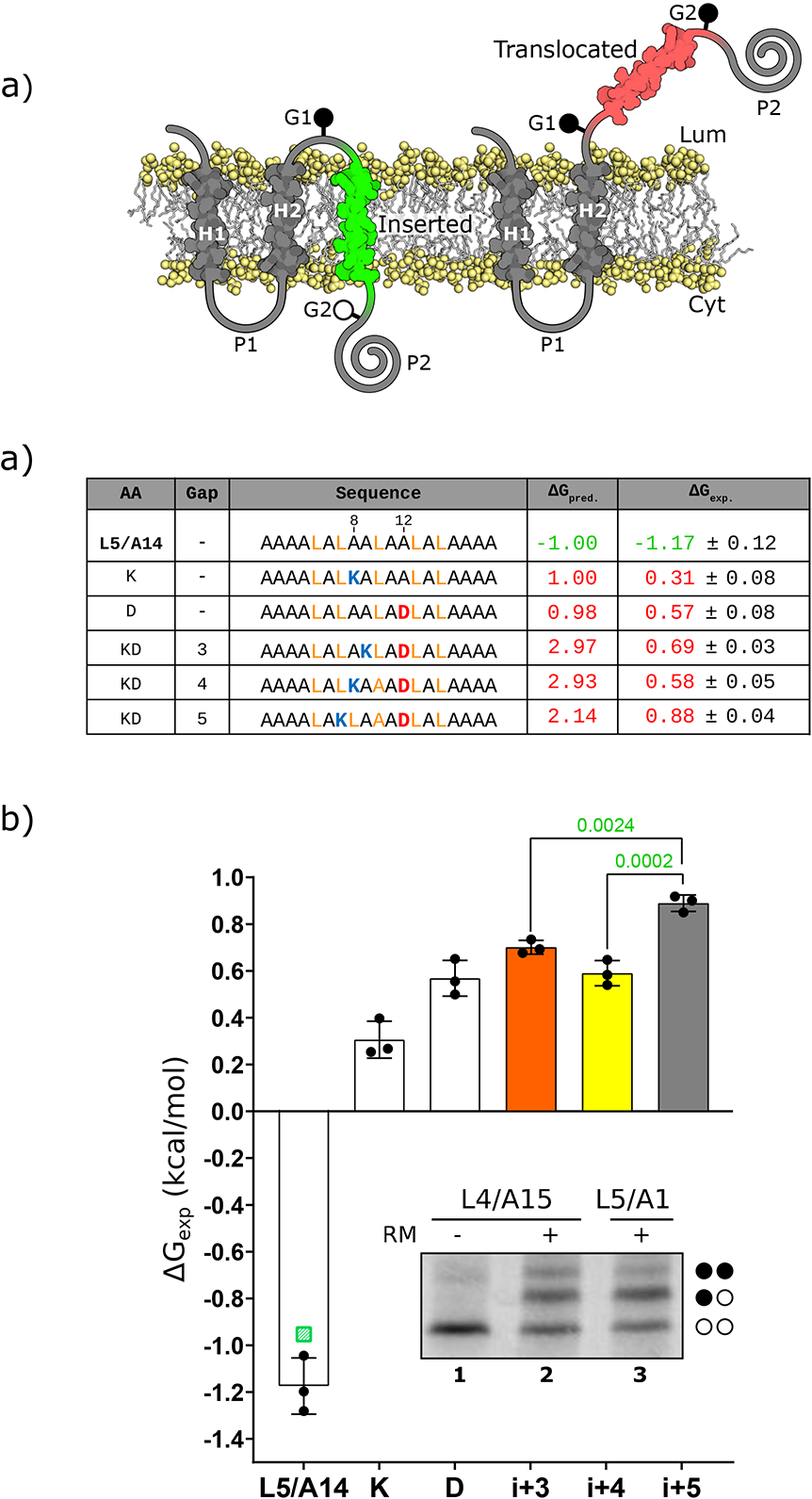
Effects on membrane insertion of single or pairs of Asp and Lys residues in L5/A14. **a)** The tested sequences from L5/A14 model TM (including the charged residues, bold), the gap distance, and the predicted ΔG (ΔG_pred_) and experimental (ΔG_exp_) values in kcal/mol are shown. Amino acids with positive and negative charge are highlighted in blue (K) and red (D) respectively. **b)** Experimental ΔG (ΔG_exp_) in kcal/mol of each tested sequence in the Lep-based microsomal assay. The mean and standard deviation of 3 independent experiments are represented. The individual value of each experiment is represented by a solid dot, *p*-values are indicated above. In addition, a green dot represents the ΔG_pred_ value for the L5/A14 sequence. The wt and single mutants are shown in white bars. Charges at compatible distances with salt bridge formation (*i*, *i*+3; and *i*, *i*+4) are shown in orange and yellow, respectively. Not compatible distances with salt bridge formation (*i*, *i*+5) is shown in gray. The inset shows a representative SDS-PAGE gel for L4/A15 and L5/A14 constructs. The construct was expressed in rabbit reticulocyte lysed in the presence (+RM) or absence (–RM) of column washed rough microsomes. Bands of non-glycosylated proteins are indicated by a white dot; mono and double glycosylated proteins are indicated by one and two black dots, respectively.

**Figure S2.**
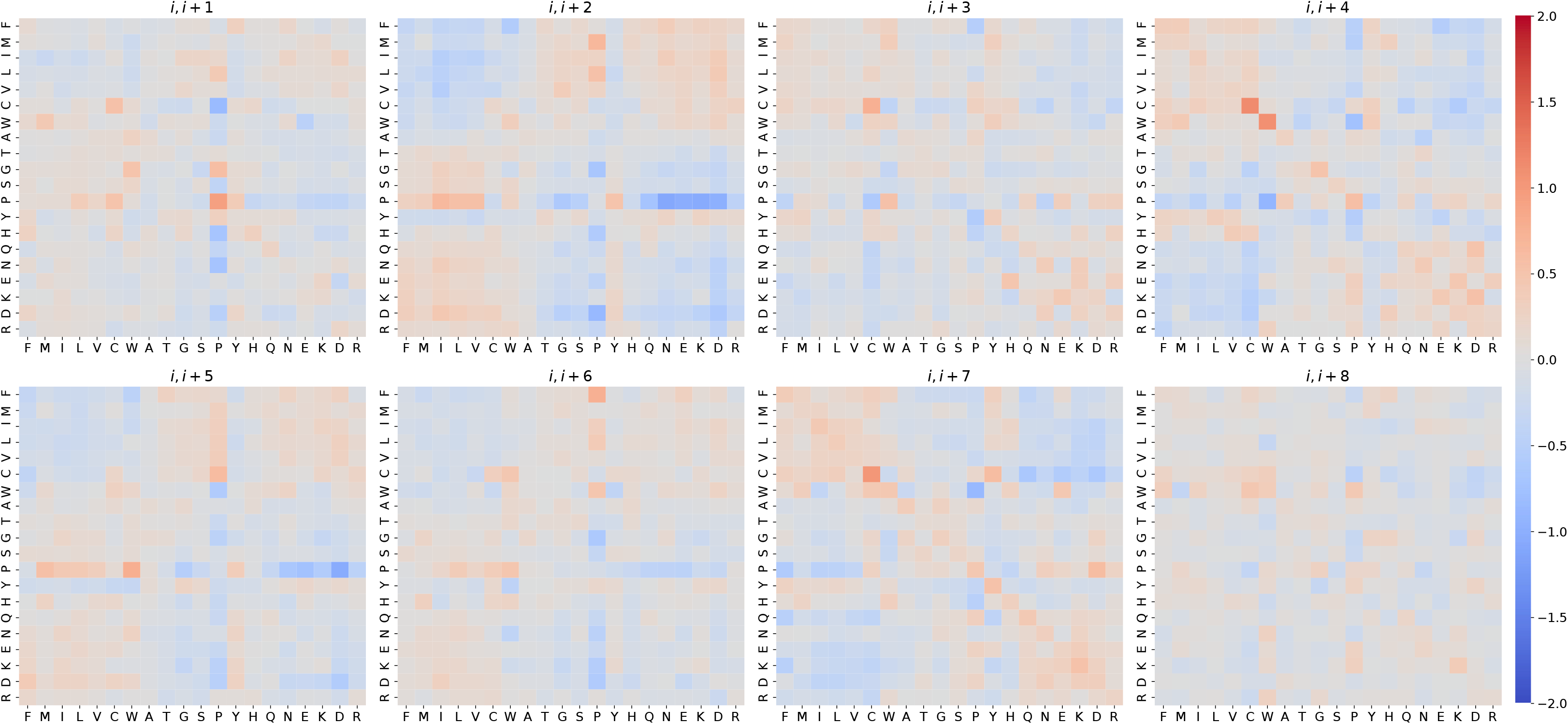
Log odds ratios of each pair of amino acids for ‘*i*, *i*+1’ through ‘*i*, *i*+8’ for the GLOB-MSA dataset. Log odds ratios for the middle core residues of α-helices of at least 17 residues in length in the GLOB-MSA dataset. The rows on the y-axis indicate the first amino acid in the pair and the columns on the x-axis the second. The residues are ordered as in Fig. 2.

**Figure S3.**
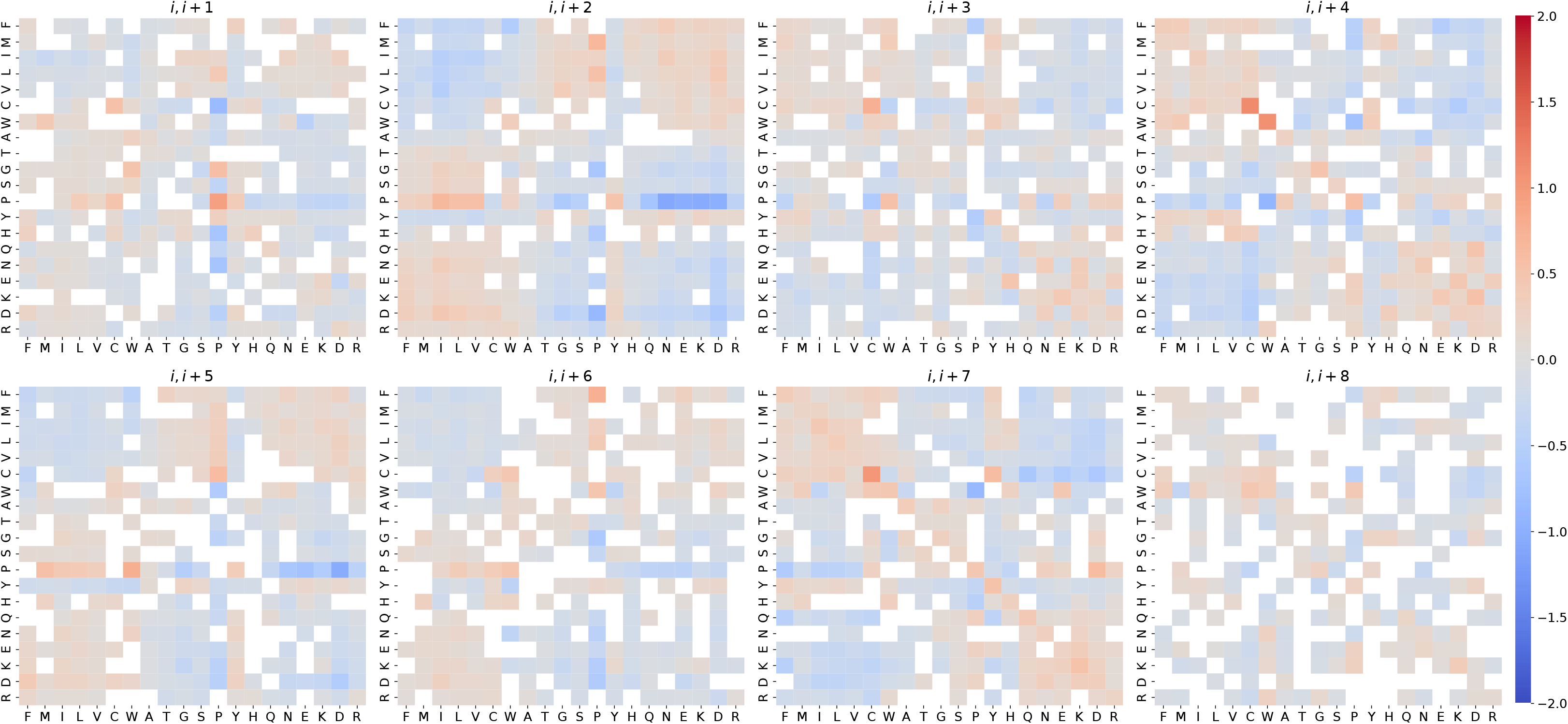
Log odds ratios of each pair of amino acids for ‘*i*, *i*+1’ through ‘*i*, *i*+8’ for the GLOB-MSA dataset. Log odds ratios as in S2 but all pairs with P-value >0.05 have been masked.

**Figure S4.**
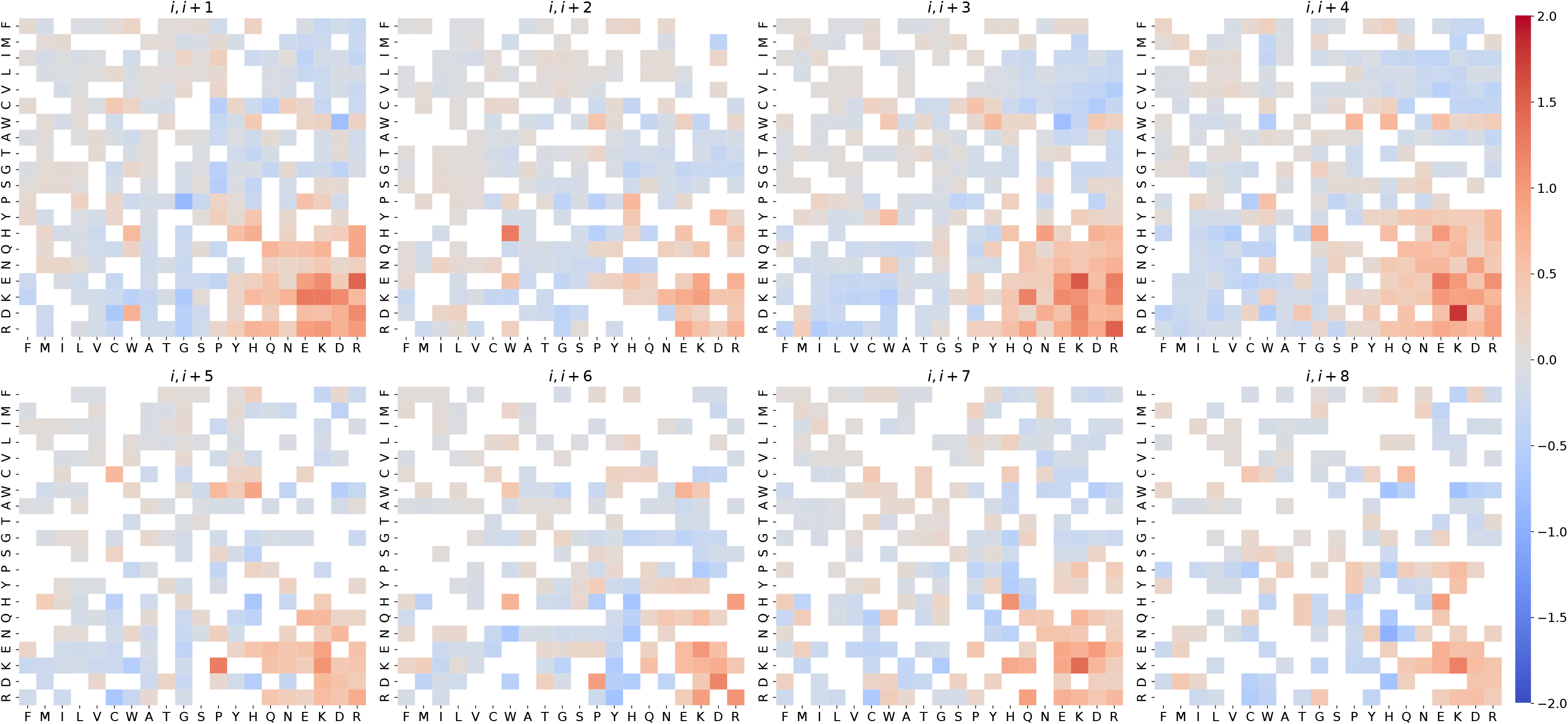
Masked log odds ratios of each pair of amino acids for ‘*i*, *i*+1’ through ‘*i*, *i*+8’ for the TM-MSA dataset. Log odds ratios as in Fig. 2 but all pairs with P-value >0.05 have been masked.

**Figure S5.**
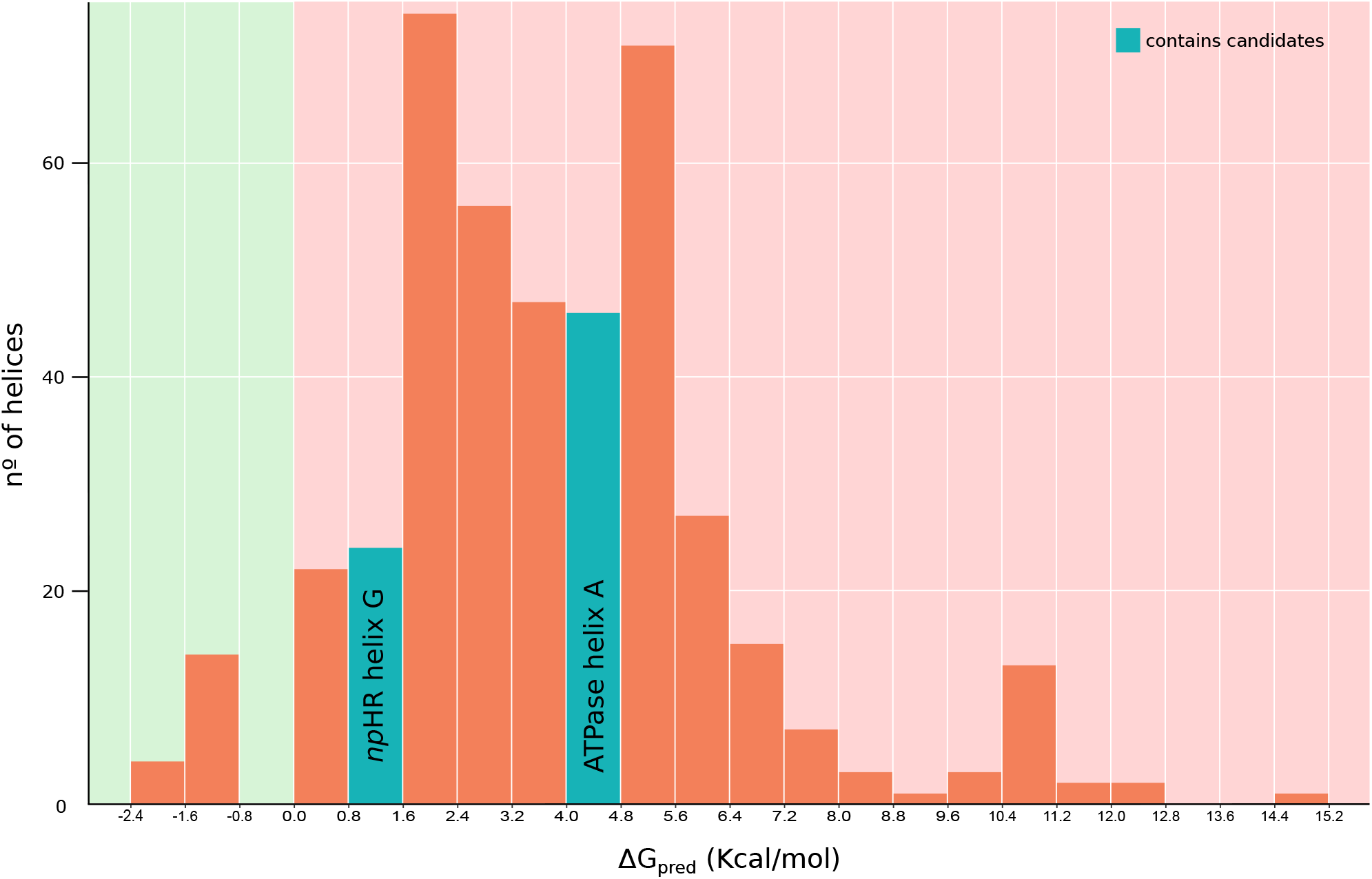
Histogram of ΔG_pred_ values for the salt bridge containing TM segments. ΔG values represented with a bin size of 0.8 kcal/mol, with negative (indicative of insertion) and positive (indicative of non-inserted) values highlighted with a green and red background, respectively. Halorhodopsin 3QBG helix G (ΔG_pred_ of 1.73 kcal/mol) and Ca2+ ATPase helix A (ΔG_pred_ of 4.15 kcal/mol) are part of the left and right blue highlighted bars, respectively.

**Figure S6.**
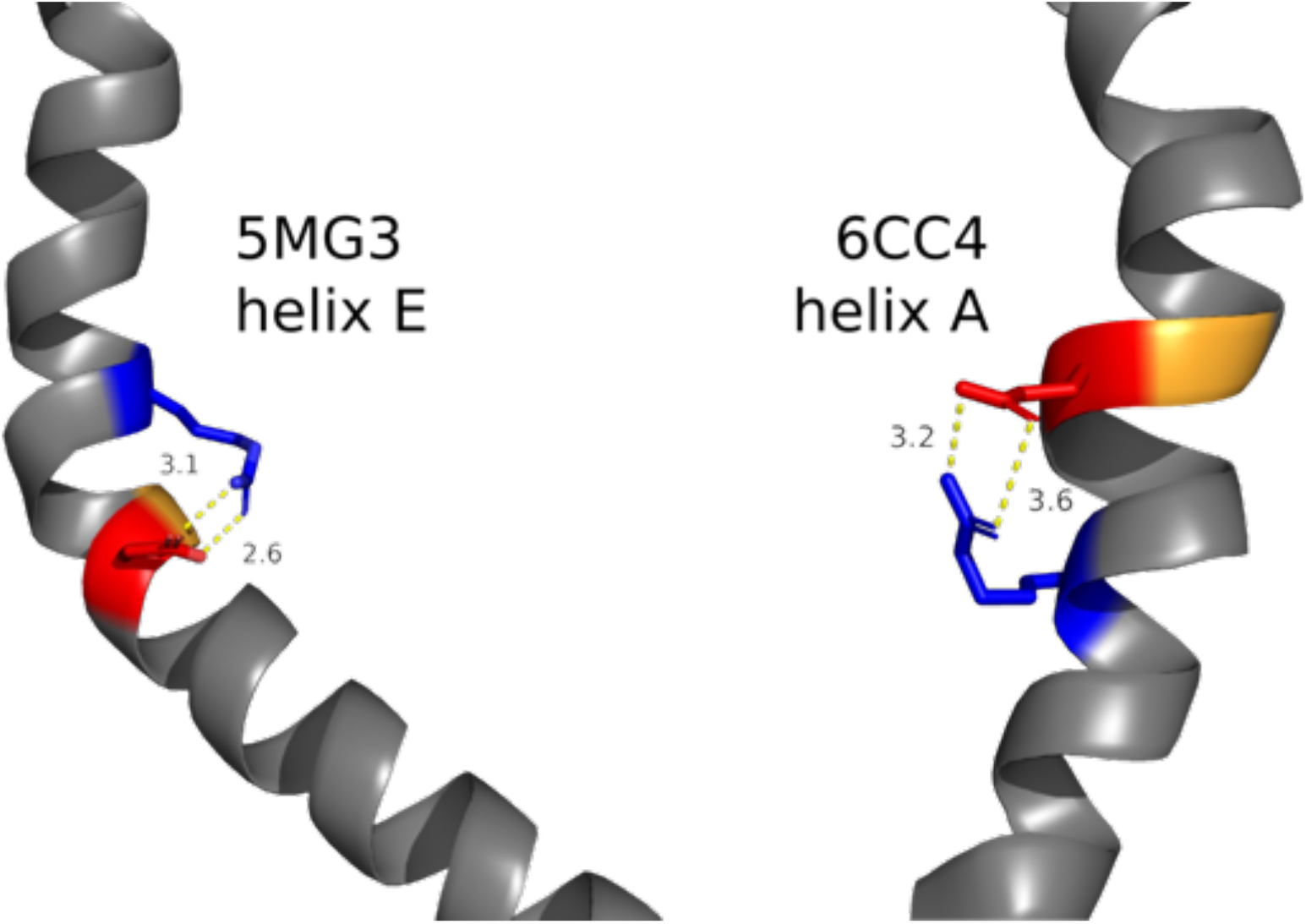
Salt bridges at pair *i*, *i*+5. Left: helix E from bacterial translocon (PDB ID: 5MG3). Glycine (orange) residue causes a kink facilitating salt bridge interaction between Asp50 (red) and Arg55 (blue). Right: helix A from a lipid flippase (PDB ID: 6CC4). Glycine (orange) residue causes a kink facilitating salt bridge interaction between Arg153 (blue) and Glu158 (red). Distances are shown in Ångström.

## REFERENCES

Andreeva, A., Howorth, D., Chothia, C., Kulesha, E., and Murzin, A.G. (2013). SCOP2 prototype: a new approach to protein structure mining. Nucleic Acids Res. 42, D310–D314.

Andreeva, A., Kulesha, E., Gough, J., and Murzin, A.G. (2019). The SCOP database in 2020: expanded classification of representative family and superfamily domains of known protein structures. Nucleic Acids Res. 48, D376– D382.

Armstrong, K.M., and Baldwin, R.L. (1993). Charged histidine affects alpha-helix stability at all positions in the helix by interacting with the backbone charges. Proc. Natl. Acad. Sci. U.S.a. 90, 11337–11340.

Baeza-Delgado, C., Heijne von, G., Marti-Renom, M.A., and Mingarro, I. (2016). Biological insertion ofcomputationally designed shorttransmembrane segments. Sci. Rep. 1–9.

Baeza-Delgado, C., Marti-Renom, M.A., and Mingarro, I. (2012). Structure-based statistical analysis of transmembrane helices. Eur Biophys J 42, 199–207.

Bañó-Polo, M., Baeza-Delgado, C., Orzáez, M., Marti-Renom, M.A., Abad, C., and Mingarro, I. (2012). Polar/Ionizable Residues in Transmembrane Segments: Effects on Helix-Helix Packing. PLoS ONE 7, e44263.

Bañó-Polo, M., Baeza-Delgado, C., Tamborero, S., Hazel, A., Grau, B., Nilsson, I., Whitley, P., Gumbart, J.C., Heijne von, G., and Mingarro, I. (2018). Transmembrane but not soluble helices fold inside the ribosome tunnel. Nat Commun 9, 5246.

Bañó-Polo, M., Martínez-Gil, L., Barrera, F.N., and Mingarro, I. (2019). Insertion of Bacteriorhodopsin Helix C Variants into Biological Membranes. ACS Omega 5, 556–560.

Bañó-Polo, M., Martínez-Gil, L., Wallner, B., Nieva, J.L., Elofsson, A., and Mingarro, I. (2013). Charge Pair Interactions in Transmembrane Helices and Turn Propensity of the Connecting Sequence Promote Helical Hairpin Insertion. J Mol Biol 425, 830–840.

Bosshard, H.R., Marti, D.N., and Jelesarov, I. (2004). Protein stabilization by salt bridges: concepts, experimental approaches and clarification of some misunderstandings. J. Mol. Recognit. 17, 1–16.

Braunger, K., Pfeffer, S., Shrimal, S., Gilmore, R., Berninghausen, O., Mandon, E.C., Becker, T., Förster, F., and Beckmann, R. (2018). Structural basis for coupling protein transport and N-glycosylation at the mammalian endoplasmic reticulum. Science 360, 215–219.

Chin, C.N., and Heijne von, G. (2000). Charge pair interactions in a model transmembrane helix in the ER membrane. J Mol Biol 303, 1–5.

Chitwood, P.J., and Hegde, R.S. (2020). An intramembrane chaperone complex facilitates membrane protein biogenesis. Nature 584, 1–22.

Donald, J.E., Kulp, D.W., and DeGrado, W.F. (2011). Salt bridges: Geometrically specific, designable interactions. Proteins 79, 898–915.

Duart, G., García-Murria, M.J., Grau, B., Acosta-Cáceres, J.M., Martínez-Gil, L., and Mingarro, I. (2020). SARS-CoV-2 envelope protein topology in eukaryotic membranes. Open Biol. 10, 200209–6.

Eddy, S.R. (2011). Accelerated Profile HMM Searches. PLoS Comput Biol 7, e1002195.

Engelman, D.M., Steitz, T.A., and Goldman, A. (1986). Identifying nonpolar transbilayer helices in amino acid sequences of membrane proteins. Annu Rev Biophys Biophys Chem 15, 321–353.

Feige, M.J., and Hendershot, L.M. (2013). Quality Control of Integral Membrane Proteins by Assembly-Dependent Membrane Integration. Mol Cell 51, 297–309.

Fu, L., Niu, B., Zhu, Z., Wu, S., and Li, W. (2012). CD-HIT: accelerated for clustering the next-generation sequencing data. Bioinformatics 28, 3150–3152.

Giménez-Andrés, M., Čopič, A., and Antonny, B. (2018). The Many Faces of Amphipathic Helices. Biomolecules 8, 45–14.

Hedin, L.E., Ojemalm, K., Bernsel, A., Hennerdal, A., Illergård, K., Enquist, K., Kauko, A., Cristobal, S., Heijne von, G., Lerch-Bader, M., et al. (2010). Membrane insertion of marginally hydrophobic transmembrane helices depends on sequence context. J Mol Biol 396, 221–229.

Heijne von, G. (1989). Control of topology and mode of assembly of a polytopic membrane protein by positively charged residues. Nature 341, 456–458.

Hessa, T., Kim, H., Bihlmaier, K., Lundin, C., Boekel, J., Andersson, H., Nilsson, I., White, S.H., and Heijne von, G. (2005). Recognition of transmembrane helices by the endoplasmic reticulum translocon. Nature 433, 377–381.

Hessa, T., Meindl-Beinker, N.M., Bernsel, A., Kim, H., Sato, Y., Lerch-Bader, M., Nilsson, I., White, S.H., and Heijne von, G. (2007). Molecular code for transmembrane-helix recognition by the Sec61 translocon. Nature 450, 1026– 1030.

Illergård, K., Kauko, A., and Elofsson, A. (2011). Why are polar residues within the membrane core evolutionary conserved? Proteins 79, 79–91.

Jayasinghe, S., Hristova, K., and White, S.H. (2001). Energetics, stability, and prediction of transmembrane helices. J Mol Biol 312, 927–934.

Kanada, S., Takeguchi, Y., Murakami, M., Ihara, K., and Kouyama, T. (2011). Crystal Structures of an O-Like Blue Form and an Anion-Free Yellow Form of pharaonis Halorhodopsin. J Mol Biol 413, 162–176.

Kauko, A., Hedin, L.E., Thebaud, E., Cristobal, S., Elofsson, A., and Heijne von, G. (2010). Repositioning of transmembrane alpha-helices during membrane protein folding. J Mol Biol 397, 190–201.

Kauko, A., Illergård, K., and Elofsson, A. (2008). Coils in the Membrane Core Are Conserved and Functionally Important. J Mol Biol 380, 170–180.

Kolbe, M., Besir, H., Essen, L.O., and Oesterhelt, D. (2000). Structure of the light-driven chloride pump halorhodopsin at 1.8 A resolution. Science 288, 1390–1396.

Kozma, D., Simon, I., and Tusnády, G.E. (2013). PDBTM: Protein Data Bank of transmembrane proteins after 8 years. Nucleic Acids Res. 41, D524–D529.

Kumar, S., Ma, B., Tsai, C.J., and Nussinov, R. (2000). Electrostatic strengths of salt bridges in thermophilic and mesophilic glutamate dehydrogenase monomers. Proteins 38, 368–383.

Kumar, S., and Nussinov, R. (2002). Relationship between Ion Pair Geometries and Electrostatic Strengths in Proteins. Biophys J 83, 1595–1612.

Lerch-Bader, M., Lundin, C., Kim, H., Nilsson, I., and Heijne von, G. (2008). Contribution of positively charged flanking residues to the insertion of transmembrane helices into the endoplasmic reticulum. 105, 4127–4132.

Li, W., and Godzik, A. (2006). Cd-hit: a fast program for clustering and comparing large sets of protein or nucleotide sequences.

Lin, C.-Y., and Lin, L.-Y. (2018). The conserved basic residues and the charged amino acid residues at the α-helix of the zinc finger motif regulate the nuclear transport activity of triple C2H2 zinc finger proteins. PLoS ONE 13, e0191971–20.

Marqusee, S., and Baldwin, R.L. (1987). Helix stabilization by Glu-…Lys+ salt bridges in short peptides of de novo design. Proc. Natl. Acad. Sci. U.S.a. 84, 8898–8902.

Martínez-Gil, L., Saurí, A., Marti-Renom, M.A., and Mingarro, I. (2011). Membrane protein integration into the endoplasmic reticulum. Febs J 278, 3846–3858.

Mbaye, M.N., Hou, Q., Basu, S., Teheux, F., Pucci, F., and Rooman, M. (2019). A comprehensive computational study of amino acid interactions in membrane proteins. Sci. Rep. 9, 12043–14.

Okamura, Y., Fujiwara, Y., and Sakata, S. (2015). Gating Mechanisms of Voltage-Gated Proton Channels. Annu Rev Biochem 84, 685–709.

Samanta, U., Pal, D., and Chakrabarti, P. (1999). Packing of aromatic rings against tryptophan residues in proteins. Acta Cryst (1999). D55, 1421-1427 [Doi:10.1107/S090744499900726X] 1–7.

Shurtleff, M.J., Itzhak, D.N., Hussmann, J.A., Schirle Oakdale, N.T., Costa, E.A., Jonikas, M., Weibezahn, J., Popova, K.D., Jan, C.H., Sinitcyn, P., et al. (2018). The ER membrane protein complex interacts cotranslationally to enable biogenesis of multipass membrane proteins. eLife 7, 382.

Tamborero, S., Vilar, M., Martínez-Gil, L., Johnson, A.E., and Mingarro, I. (2011). Membrane insertion and topology of the translocating chain-associating membrane protein (TRAM). J Mol Biol 406, 571–582.

Toyoshima, C., Nakasako, M., Nomura, H., and Ogawa, H. (2000). Crystal structure of the calcium pump of sarcoplasmic reticulum at 2.6 Å resolution. Nature 405, 647–655.

Tsirigos, K.D., Govindarajan, S., Bassot, C., Västermark, Å., Lamb, J., Shu, N., and Elofsson, A. (2018). ScienceDirect Topology of membrane proteins — predictions, limitations and variations. Curr Opin Struct Biol 50, 9–17.

Tsirigos, K.D., Peters, C., Shu, N., Käll, L., and Elofsson, A. (2015). The TOPCONS web server for consensus prediction of membrane protein topology and signal peptides. Nucleic Acids Res. 43, W401–W407.

Walther, T.H., and Ulrich, A.S. (2014). Transmembrane helix assembly and the role of salt bridges. Curr Opin Struct Biol 27, 63–68.

